# HIV rapidly targets a diverse pool of CD4+ T cells to establish productive and latent infections

**DOI:** 10.1101/2022.05.10.491275

**Authors:** Pierre Gantner, Supranee Buranapraditkun, Amélie Pagliuzza, Caroline Dufour, Marion Pardons, Julie L. Mitchell, Eugène Kroon, Carlo Sacdalan, Nicha Tulmethakaan, Suteeraporn Pinyakorn, Merlin L. Robb, Nittaya Phanuphak, Jintanat Ananworanich, Denise Hsu, Sandhya Vasan, Lydie Trautmann, Rémi Fromentin, Nicolas Chomont

**Author notes:** Corresponding author: Nicolas Chomont, Centre de Recherche du CHUM, 900 rue St-Denis, Montreal, H2X 0A9, QC, Canada; #31266.

## Abstract

Upon infection, HIV disseminates throughout the human body within 1-2 weeks. However, its early cellular targets remain poorly characterized. We analyzed productively and latently infected cells in blood and lymphoid tissue from individuals in acute infection. The phenotype of productively infected cells rapidly evolved with time and differed between blood and lymph nodes. The TCR repertoire of productively infected cells was heavily biased, with preferential infection of previously expanded/disseminated cells, but composed almost exclusively of unique clonotypes, indicating that they were the product of independent infection events. Latent genetically intact proviruses were already archived early in infection. Hence, productive infection is initially established in a pool of phenotypically and clonotypically distinct T cells in blood and lymph nodes and latently infected cells are generated simultaneously.

**One-Sentence Summary:** HIV initially infects phenotypically and clonotypically distinct T cells and establishes a latent reservoir concomitantly.

## Main Text

Studying acute HIV is key to understand initial infection events and to identify the mechanisms leading to the establishment of the persistent viral reservoir. More than 40 years after its discovery (*1*), the early cellular targets of HIV within the first days/weeks of infection remain uncharacterized. Since accessing lymphoid tissues in people who recently acquired HIV is difficult, viral dissemination is hard to study in humans and most observations have been made in SIV-infected non-human primate models: Upon mucosal challenge, SIV rapidly establishes a small pool of productively infected cells that disseminate within days to lymphoid tissues and peripheral blood (*2*), following local replication of transmitted founders at mucosal sites (*3, 4*). This early phase of SIV infection is characterized by a profound depletion of CD4+ T cells in blood and tissues (*5*), particularly in the intestinal tract (*6*). During chronic infection, the bulk of SIV replication is thought to occur in gut and lymph nodes (*7*). However, several parameters differ between non-human primate models and HIV-infected individuals (e.g. viruses, routes of infection, inoculum), hence the mechanisms of HIV dissemination in humans remain elusive.

In humans, we previously reported that maximal levels of HIV DNA are reached in lymph nodes as early as Fiebig stages II/III (*8*). Although it is well established that T follicular helper cells (Tfh) are preferential targets for HIV in chronic infection (*9*), their contribution to the early dissemination of HIV is currently unknown. While productively infected cells usually do not survive for more than 2 days (*10, 11*), they are rapidly replaced and maintain the rate of new infections which causes a rapid rise in plasma viremia during the first few weeks of infection (*12*). Little is known about the nature and the location of the cells in which HIV establishes productive infection in lymphoid tissues and blood during the initial phase of infection (i.e. before seroconversion, Fiebig stages I and II), and how these cells contribute to viral dissemination. Although duplications of HIV integration sites within the pool of HIV DNA-bearing cells are rarely detected during late stages of acute infection (*13, 14*), it remains unclear whether antigen-driven expansions (particularly those resulting from HIV antigens) contribute to the initial dissemination of productively infected cells.

HIV reservoirs are established in the form of latently infected CD4+ T cells during acute infection (*2, 15*) and persist for decades on ART through clonal expansion (*16–19*). The cellular and molecular mechanisms contributing to the establishment of HIV latency *in vivo* are still poorly elucidated. *In vitro*, activated CD4+ T cells transitioning to resting state in a relatively narrow time window are highly permissive for latent infection of R5-tropic HIV (*20*). An alternative and non-exclusive mechanism for the establishment of HIV latency is provided by the ability of specific chemokines to increase the permissiveness of resting CD4+ T cells to HIV infection up to the proviral integration step (*21, 22*). The concept that latency is established during acute infection essentially stems from observational studies showing that viral rebound is observed upon treatment interruption even when ART is initiated during acute HIV infection (*2, 15, 23*). However, whether genetically intact genomes are already archived in the latent reservoir at this early stage has not yet been reported.

We used a combination of single-cell approaches in blood and lymph nodes from individuals at the earliest stages of HIV infection to investigate the location and mechanisms by which productive infection is established in acute infection. In addition, we obtained near full-length HIV genomes from longitudinal samples of early treated individuals to characterize the fate, inducibility, and genetic intactness of proviruses archived early in infection.

## Results

### Productively infected cells are detected in blood and lymph nodes from the earliest stages of acute HIV infection

The RV254/SEARCH 010 cohort enrolls acutely infected individuals in Fiebig stages I-V who initiate ART within a median time of 2 days after diagnosis in Bangkok, Thailand (*8, 24*). To quantify and characterize the initial pool of productively infected cells in blood and tissues, leukapheresis and inguinal lymph nodes biopsies were obtained at the time of diagnosis in acute infection prior to ART initiation in a subset of consenting participants. Paired PBMCs (peripheral blood mononuclear cells) and LNMCs (lymph node mononuclear cells) from 21 acutely infected participants of the RV254/SEARCH 010 study were obtained (Table S1). PBMCs and LNMCs from 4 chronically infected and ART-naïve Thai individuals enrolled in the RV304/SEARCH 013 study were also obtained. To measure the frequency of productively infected cells and assess their phenotype, we used HIV-Flow (Fig. S1) (*25*), which captures cells expressing the p24 capsid protein in enriched CD4+ T cells. Productively infected cells were readily detected in blood and lymph node samples from all participants (Fig. 1a), including in all 5 individuals at Fiebig stage I (11 to 16 days post infection (*26*), median frequencies = 3.1 and 4.5 p24+ cells / million cells in blood and lymph nodes, respectively). Median frequencies of productively infected cells increased during the transition to Fiebig stage II, reached maximal values at Fiebig stage III (medians, 1,100 and 100 p24+ cells / million cells in blood and lymph nodes, respectively) and remained relatively stable afterwards, including during chronic infection. Frequencies of p24+ cells strongly correlated between blood and lymph nodes (Fig. 1b) and the frequency of p24+ measured in the blood, but not in the lymph nodes, correlated with plasma viremia (Fig. 1c). These results indicate that the frequency of productively infected cells rapidly increases during acute infection and that up to 1/1,000 circulating CD4+ T cells is productively infected when plasma viremia peaks (Fiebig stage III).

**Fig. 1.**
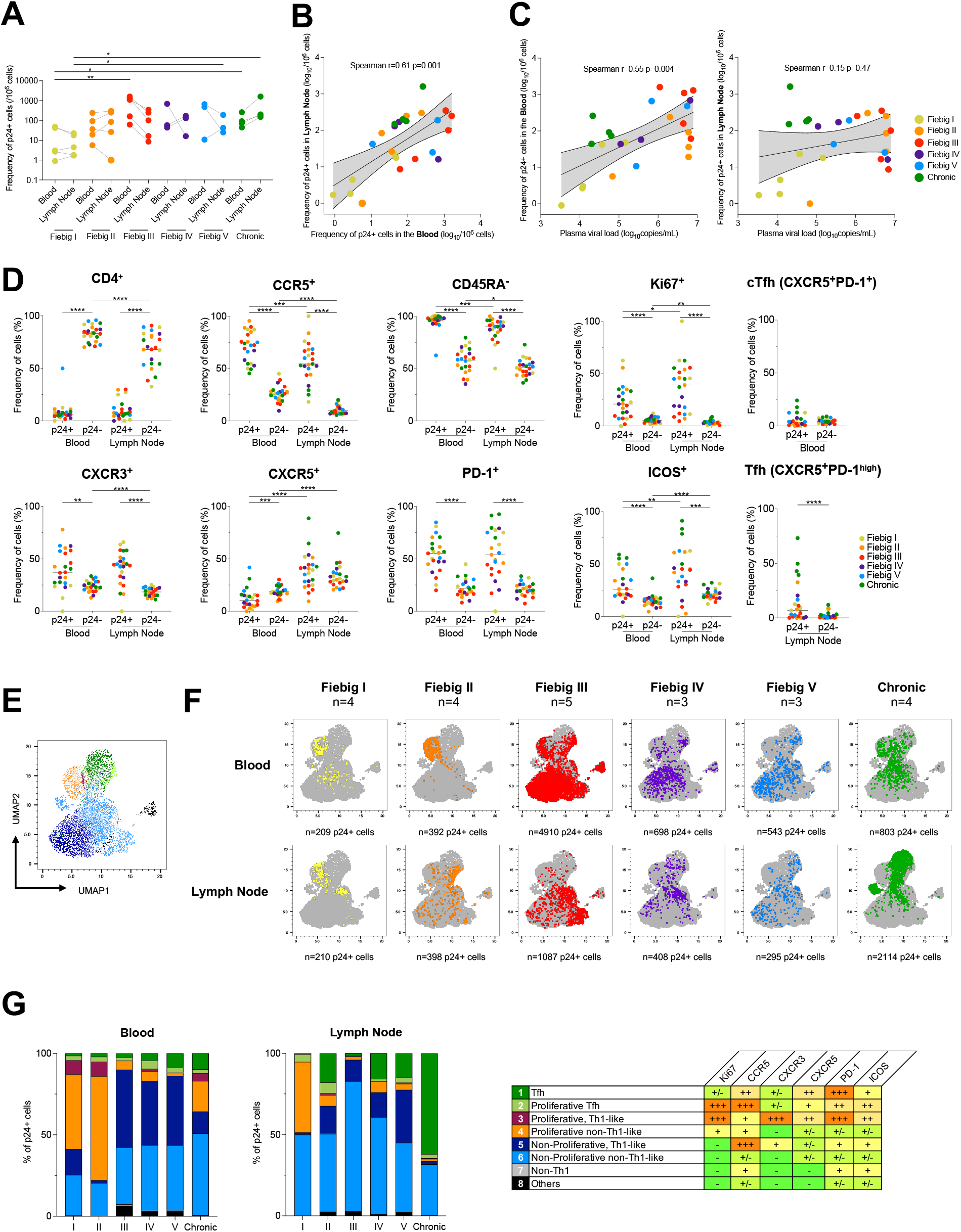
Frequencies and phenotype of productively infected cells. **A.** Frequencies of p24+ cells in CD4+ T cells measured by HIV-Flow in paired blood and lymph node samples from participants at different Fiebig stages of acute infection and in chronically infected controls. **B.** Correlation between p24+ cells frequencies in blood and lymph nodes. **C.** Correlations between p24+ cells frequencies in blood or lymph nodes with plasma viremia. **D.** Phenotype of productively infected cells. Frequencies of p24+ and p24-cells from blood and lymph node expressing each marker or combination of markers (CD4-, CCR5+, CD45RA-, Ki67+, CXCR3+, CXCR5+, PD-1+, ICOS+, circulating T follicular helpers (cTfh) cells and Tfh cells) are depicted for each participant. **E.** p24+ cells phenotypic data were integrated in a UMAP analysis and generated 8 cell clusters. **F.** Dot plots of the UMAP analysis of all p24+ cells (in grey) as shown in panel e, on which the p24+ cells by stage of infection and compartment are overlaid in colors. **G.** The frequency of each of these clusters among p24+ cells is depicted in bar graphs according to the stage of infection for both blood and lymph node. The table shows the phenotype of each cell cluster. Statistically significant differences are highlighted (Wilcoxon; p<0.05, *; p<0.01**; p<0.001, ***; p<0.0001 ****).

### Several cellular markers are preferentially expressed by productively infected cells from the earliest stages of acute infection

To identify markers specifically expressed by productively infected cells, we analyzed the phenotypes of p24+ cells in blood and lymph nodes from all participants (Fig S2). As expected, most p24+ cells expressed low levels of the CD4 receptor when compared to their uninfected or infected but translation-incompetent counterparts (Fig. 1d), which likely resulted from its downregulation from the cell surface by nef, vpu and env (*27*). Productively infected cells from both blood and lymph nodes frequently expressed CCR5 (the major HIV-co-receptor), displayed a memory phenotype (CD45RA-), and were enriched in subsets of activated (Ki67+, ICOS+, PD-1+) and Th1-like (CXCR3+) cells. The phenotypic signature of productively infected cells changed over time: whereas the contribution of lymph node T follicular helper cells (Tfh, CXCR5+PD-1^high^) to the initial pool of infected cells was modest at all stages of acute infection (<15%, Fig. 1d and Fig S3a), Tfh cells were major HIV producers in lymph nodes from chronically infected participants (*9*) and encompassed 45% of all productively infected cells. Over time, p24+ producing cells tended to express less frequently CCR5 and more frequently ICOS (Fig. S3a), although interindividual variations were large. When compared to their uninfected counterparts, the initial pool of infected cells in the lymph nodes from Fiebig I participants were 40 times more likely to express Ki67 and 7 times more likely to express CCR5 (Fig. S3b). Altogether, these observations indicate that during acute infection, productively infected cells are primarily memory Th1-like T cells that do not express CD4 at the cell surface, display moderate to high levels of activation markers and the CCR5 HIV co-receptor, and are unlikely to exert Tfh functions.

### The phenotype of infected cells changes over time and varies between blood and lymph nodes

We next sought to compare the phenotype of p24+ cells between blood and lymph nodes at all stages of HIV infection. A UMAP analysis revealed that infected cells were distributed in 8 cell clusters (Fig. 1e and Fig. S4a). The relative contribution of each cluster to the overall pool of infected cells evolved rapidly during acute infection and differed between blood and lymph nodes (Fig. 1f). Proliferating T-cells (Ki67+, clusters 2, 3 and 4) were the major contributors to the pool of productively infected cells in both blood and lymph node during early Fiebig stages (I and II). There was a rapid shift to non-proliferating cells (cluster 5 and 6) in both blood and lymph node, some of which expressed high levels of CCR5 (cluster 5), especially in blood (Fiebig III, IV, V; Fig. 1g and Fig. S4b). During chronic infection, Tfh (cluster 1) emerged as the major contributor to the pool of infected cells in the lymph node, representing an average of 62% of all productively infected cells, while the phenotypes of p24+ cells in the blood were more diverse. Altogether, these results indicate that the cell subsets that support productive HIV infection rapidly evolve throughout acute HIV infection, differ between blood and lymph nodes, and that in contrast to chronic infection, Tfh cells marginally contribute to the initial dissemination of HIV.

### Proviral diversity is limited in productively infected cells during acute infection

We then investigated if the diversity observed in the phenotype of productively infected cells was accompanied by genetic diversity in the proviral populations. We obtained *env* (C2-V5) sequences from 534 single HIV-infected cells from n=7 participants representing all stages of infection (Fig. 2a). As expected in recently infected participants, the majority of p24+ cells harbored a single C2-V5 *env* sequence (all CCR5-tropic) that was shared between blood and lymph nodes (mean, 87.4%, Fig. 2b). In sharp contrast, only a small proportion of p24+ cells isolated during chronic infection harbored identical proviral sequences (6.2%), reflecting the diversification of viral populations during untreated progressive HIV infection. Altogether, these results suggest that a small number of viral quasispecies initially infects a small pool of phenotypically diverse cells during acute HIV infection.

**Fig. 2.**
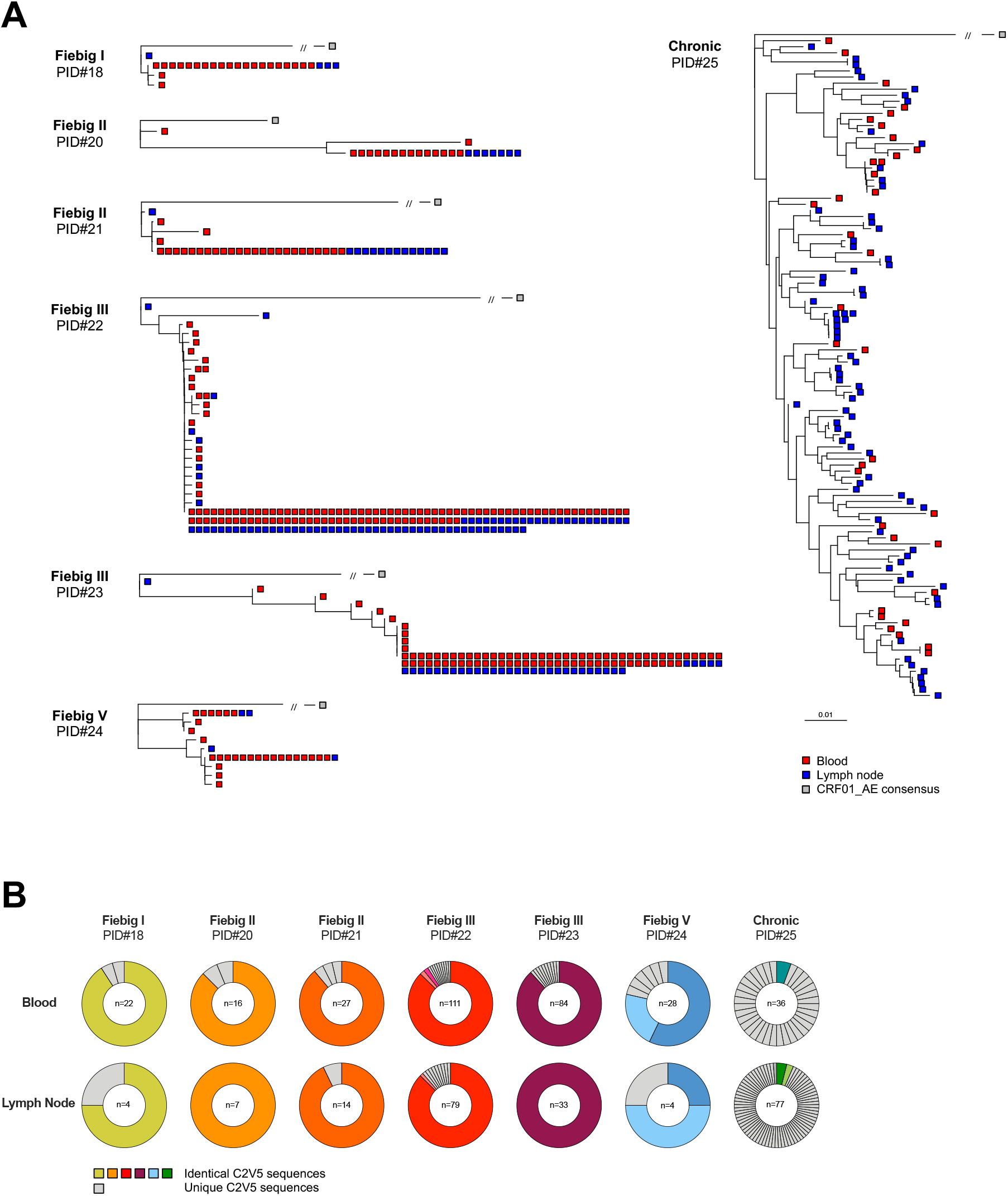
Proviral *envelope* sequence diversity in productively infected cells. **A.** Phylogenetic trees representing HIV C2-V5 *env* proviral sequences obtained from single-sorted p24+ cells from n=7 participants (PID#18 to #25). Trees were rooted on a CRF01_AE consensus (grey square) and *env* sequences from each single p24+ cell are depicted in red or blue according to their compartments (blood or lymph node, respectively). Identical sequences are shown as aggregated on the same branch of the tree. **B.** Pie charts representing the relative proportion of each C2V5 *env* sequence for each participant in both blood and lymph node. The number of p24+ cells analyzed is indicated in the center of the pie. Identical sequences are depicted in colors, whereas unique sequences are depicted in grey.

### Productive HIV infection is established in clonotypically distinct T cells from the earliest stages of HIV infection

To determine if the limited proviral diversity observed during acute infection was attributed to clonal expansions of “founder” infected T cells or to independent infection events with identical HIV variants, we combined single-cell sorting of HIV-infected cells with sequencing of the V-J junction of the TCRβ chain, which can be used as a measure of T cell clonality (*16*). Clonotypes obtained from 1,048 single HIV-infected cells isolated from 17 participants were defined either as expanded (i.e. detected in at least two cells) or unique (i.e. detected in no more than one cell). In sharp contrast to the restricted viral diversity described above, unique TCRβ clonotypes were retrieved in the overwhelming majority of productively infected cells (median, >99%), at all stages of infection and both in blood and lymph nodes (Fig. 3a). Small clonal expansions of productively infected cells (encompassing 2 to 12 p24+ cells) were detected in 6/17 participants (PID# 6, 8, 11, 12, 15 and 16), with an overall mean proportion of clonally expanded p24+ cells of 2.2% in the blood (95% CI, 0-5.1) and 0.6% in the lymph node (95% CI, 0-1.6, Fig. 3b). Of note, shared p24+ clonotype between blood and lymph nodes were not observed in any participant. In an attempt to predict if these highly diverse clonotypes could be specific to common antigens, we compared our sequences to a CDR3 database of TCR sequences with known specificities (*16, 28*). Various antigen specificities were inferred for a small fraction (8%) of p24+ clonotypes (Fig. 3c), with no obvious temporal association (Fig. 3d). Interestingly, two participants at Fiebig stage III displayed p24+ clonotypes with different CDR3 sequences sharing a same *M. tuberculosis* antigenic specificity in both blood and lymph node (Fig. 3e), suggesting that *M. tuberculosis-specific* cells may contribute to HIV dissemination in these participants. However, *M. tuberculosis-specific* cells were underrepresented in the pool of p24+ cells when compared to total CD4+ T cells (Fig. 3c). Altogether, these data indicate that HIV-infection is established by a limited number of HIV variants infecting a large pool of clonotypically distinct T cells in both blood and lymph nodes and suggest that CD4+ T cells specific to common antigens or vaccines may constitute early targets for HIV during acute infection.

**Fig. 3.**
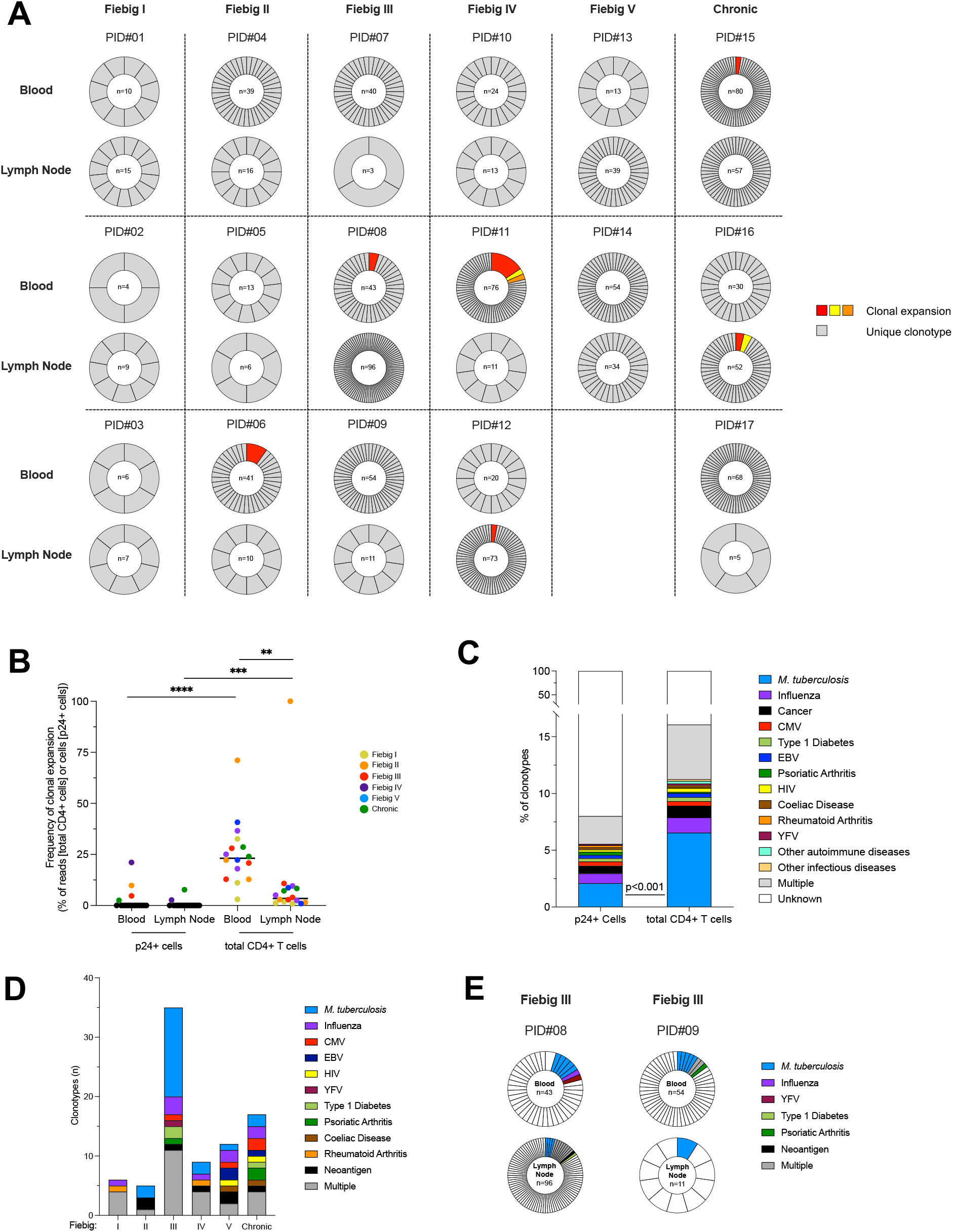
TCRβ clonotypes and clonal expansion in the pool of productively infected cells. **A.** Frequencies of TCRβ clonotypes in p24+ cells are represented for n=17 participants (PID#01 to #17) and ordered according to the stage of acute infection (Fiebig I to V) followed by chronic controls (columns) and according to the compartment (i.e. blood and lymph node, rows). For each sample, the proportion of each clonotype in the pool of p24+ cells is represented in a pie chart. The number of p24+ cells analyzed is indicated in the center of the pie. Expanded clonotypes are depicted in colors; Unique clonotypes are depicted in grey. **B.** The proportion of clonal expansions in the pool of p24+ or total CD4+ T cells is depicted for each participant. Frequencies were calculated as follows: (1) for p24+ cells, the proportion of expanded cells within the pool of p24+ cells; and (2) for total CD4+ T cells, the number of reads of expanded cells within the total number of reads in CD4+ T cells. **C.** The frequency of predicted antigen specificities is represented in the pool of p24+ or total CD4+ T clonotypes. The frequency of *M. tuberculosis* specific clonotypes was lower in p24+ cells than in total CD4+ T cells (Fisher’exact Test; p<0.001). **D.** The number of predicted antigen specificities for p24+ clonotypes is represented according to the stage of infection. **E.** Example of two participants who harbored distinct p24+ clonotypes with common antigenicity (*M. tuberculosis*). Significant differences are highlighted (Wilcoxon; p<0.05, *; p<0.01**; p<0.001, ***; p<0.0001 ****).

### HIV preferentially infects cells expressing specific Vβ and Jβ motifs

To identify potential biases in the TCR repertoire of productively infected cells, we compared the TRBV and TRBJ usage in p24+ cells with the global CD4+ T cell repertoire from the same participants. To achieve this, we applied the same TCRβ chain PCR amplification approach to 100,000 blood and lymph node CD4+ T cells from 16 participants, followed by MiSeq sequencing and analyzed by MiXCR (*29*). There was no systematic bias in the total numbers of reads and clonotypes between blood and lymph node, allowing us to compare the repertoires between these two compartments (Fig. S5a). TRBV and TRBJ usage in total CD4+ T cells were highly conserved between participants and across compartments, whereas they were more variable in p24+ cells, probably due to lower number of cells analyzed (Fig. S5b). TRBV3, 4 and 28 families and TRBJ2-1, 2-5 and 2-7 were more frequently used by p24+ cells compared to total CD4+ T cells (Fig. 4a and Fig. S5c). Conversely, TRBV12, 18, 19, 29 and 30 families and TRBJ1-2, 1-6 and 2-6 were underrepresented in p24+ cells. Collectively, these results indicate that the TCR repertoire of productively infected cells is biased when compared to the repertoire of all CD4+ T cells (Fig. S6).

**Fig. 4.**
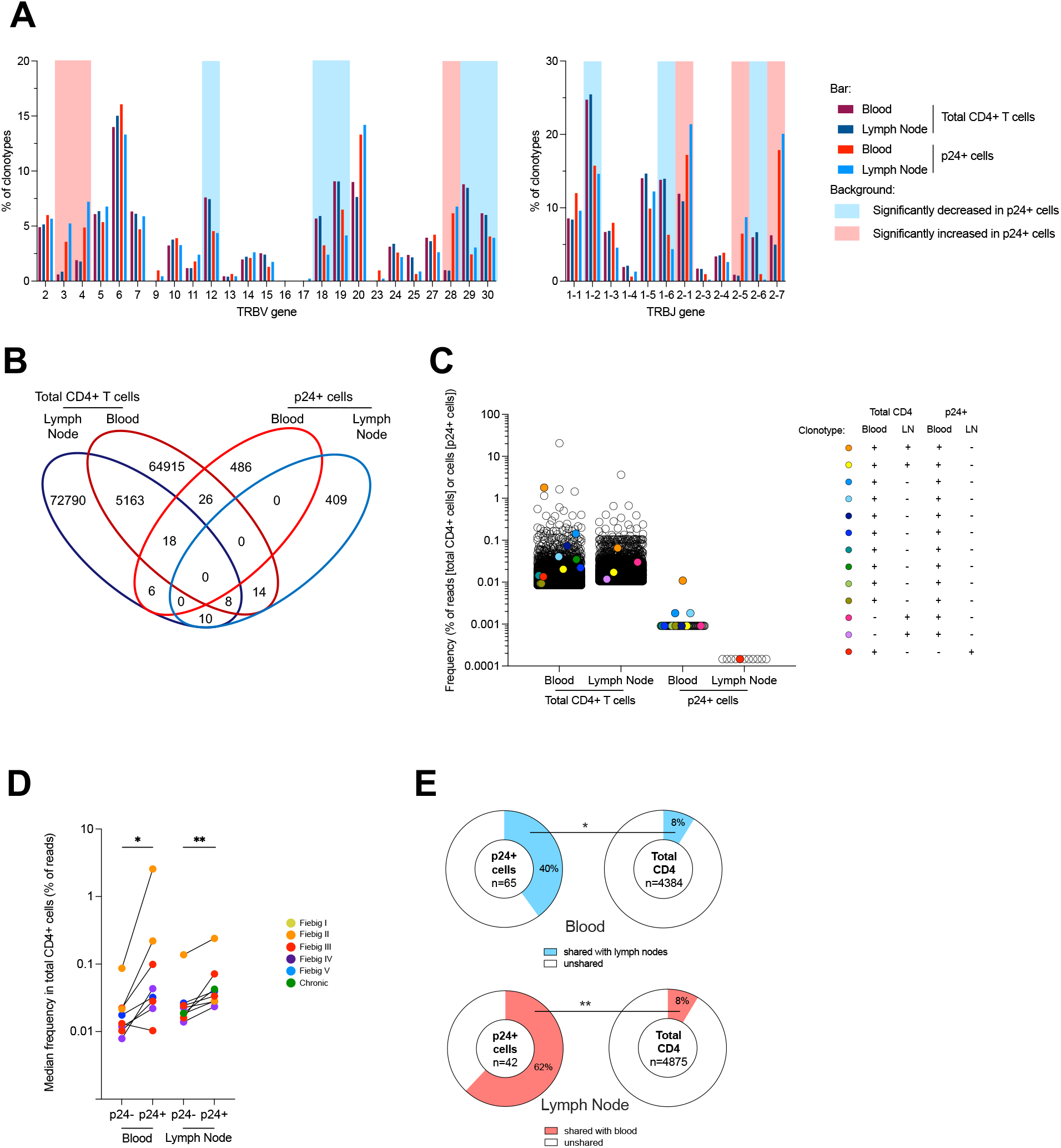
Comparison of the TCRβ repertoire of productively infected cells and total CD4+ T cells. **A.** Frequency of TRBV and TRBJ segment usage for the clonotypes identified by TCRβ sequencing in p24+ cells and in total CD4+ cells in both blood and lymph nodes. Significant differences between p24+ and total CD4+ T cells are highlighted with a red or blue background depending on the trend (see Fig. S5c). **B.** Venn diagrams showing the number of unique and shared clonotypes in the four subsets: blood and lymph node total CD4+ T cells and p24+ cells for all participants **C.** Example (Participant PID#11, data from other participants are shown in Fig. S7d) of frequency distribution (based on bulk deep sequencing data and single-cell sorting/Sanger sequencing) of the clonotypes corresponding to the clones that were found in a single subset (empty circles) and in multiple subsets including p24+ cells (colored circles). Frequencies are shown as percentage of total reads for total CD4+ T cells and as the frequency of cells determined by HIV-Flow for p24+ cells. **D.** Median frequency of reads of CD4+ T cells clonotypes from blood and lymph node in which p24+ cells were identified (p24+) or not (p24-) (data from n=9 participants with at least n=3 p24+ clonotypes shared with blood or lymph nodes are shown). **E.** Pie charts representing the median frequency of p24+ cells and total CD4+ T cells clonotypes from a given compartment (blood or lymph node) that were shared with the other compartment (lymph node or blood), respectively. The total number of cells (p24+ cells) and median number of reads (total CD4+ T cells) per condition is indicated at the center of the pie. Significant differences are highlighted (Wilcoxon or Fisher’exact test; p<0.05, *; p<0.01**; p<0.001, ***; p<0.0001 ****).

### Productive HIV infection is preferentially established in previously expanded and disseminated clonotypes

We then examined the distribution of the TCR repertoires and searched for shared clonotypes between p24+ and total CD4+ T cells. The distribution of the CD4+ T cell repertoire revealed that expanded clones were usually larger in the blood than in lymph nodes (Fig. S7a and S7b). Shared clonotypes were observed between compartments, accounting for approximately 8% of clonotypes in both compartments, and were also more expanded in the blood (median, 23% of reads) than in the lymph node (median, 11% of reads, p=0.0017, Fig. S7c). Among those expanded T cell clones shared between blood and lymph node, 82 clonotypes contained p24+ cells and were identified in 14/16 participants (Fig. 4b). Fig. 4c shows data from a representative example (participant #11): 13 clonotypes retrieved from p24+ cells were also detected in total CD4+ T cells (data from the other participants are presented in Fig. S7d). Interestingly, the T cell clones in which p24+ cells were detected were significantly larger than those in which p24+ were not retrieved (Fig. 4d), indicating that productive infection is preferentially established in individual CD4+ T cells belonging to relatively large T cell clones. In addition, the CD4+ T cells clonotypes in which p24+ cells were detected were more likely to be shared between blood and lymph nodes (Fig. 4e). Collectively, these observations indicate that previously expanded and disseminated CD4+ T cell clones are preferential targets for productive HIV infection.

### Intact and non-inducible proviruses are established during acute infection

Having demonstrated that productive HIV infection is rapidly established in phenotypically and clonotypically diverse CD4+ T cells, we sought to determine if latently infected cells could also be generated during the acute phase of infection. We used longitudinal blood samples from individuals initiating ART during acute (n=6) or chronic (n=2) infection collected before ART initiation and after 96 weeks of suppressive ART. Isolated CD4+ T cells were rested or stimulated with PMA/ionomycin to induce viral gene and protein expressions. In samples collected before ART, stimulation did not significantly increase the frequency of cells producing the viral protein p24 (Fig. 5a), nor the amount of cell-associated LTR-gag or tat/rev transcripts (Fig. 5b and Fig. S8a), indicating that latent and inducible proviruses were rare during untreated infection. To determine if productively infected cells display cellular features that may favor their long-term persistence during ART, we analyzed their phenotype (Fig. S9). Most p24+ cells displayed a central memory (T_CM_, mean, 28.3%) or an effector memory (T_EM_, mean, 50.4%) phenotype (Fig. S8b) and were in a resting state as shown by low to moderate levels of expression of the activation markers CD69 and HLA-DR (median, 7.1% and 21.0%, respectively; Fig. S8c). In addition, productively infected cells displayed lower levels of HLA class I and similar levels of the prosurvival molecule Bcl-2 than their uninfected counterparts, suggesting potential escape/survival mechanisms (Fig. S8d). Nonetheless, after 96 weeks of ART, none of the participants who initiated ART during acute infection displayed detectable translation-competent reservoir cells, even after stimulation (Fig. 5a), which was consistent with low to undetectable levels of cell-associated viral transcripts in these samples (Fig. 5b and Fig. S8a).

**Fig. 5.**
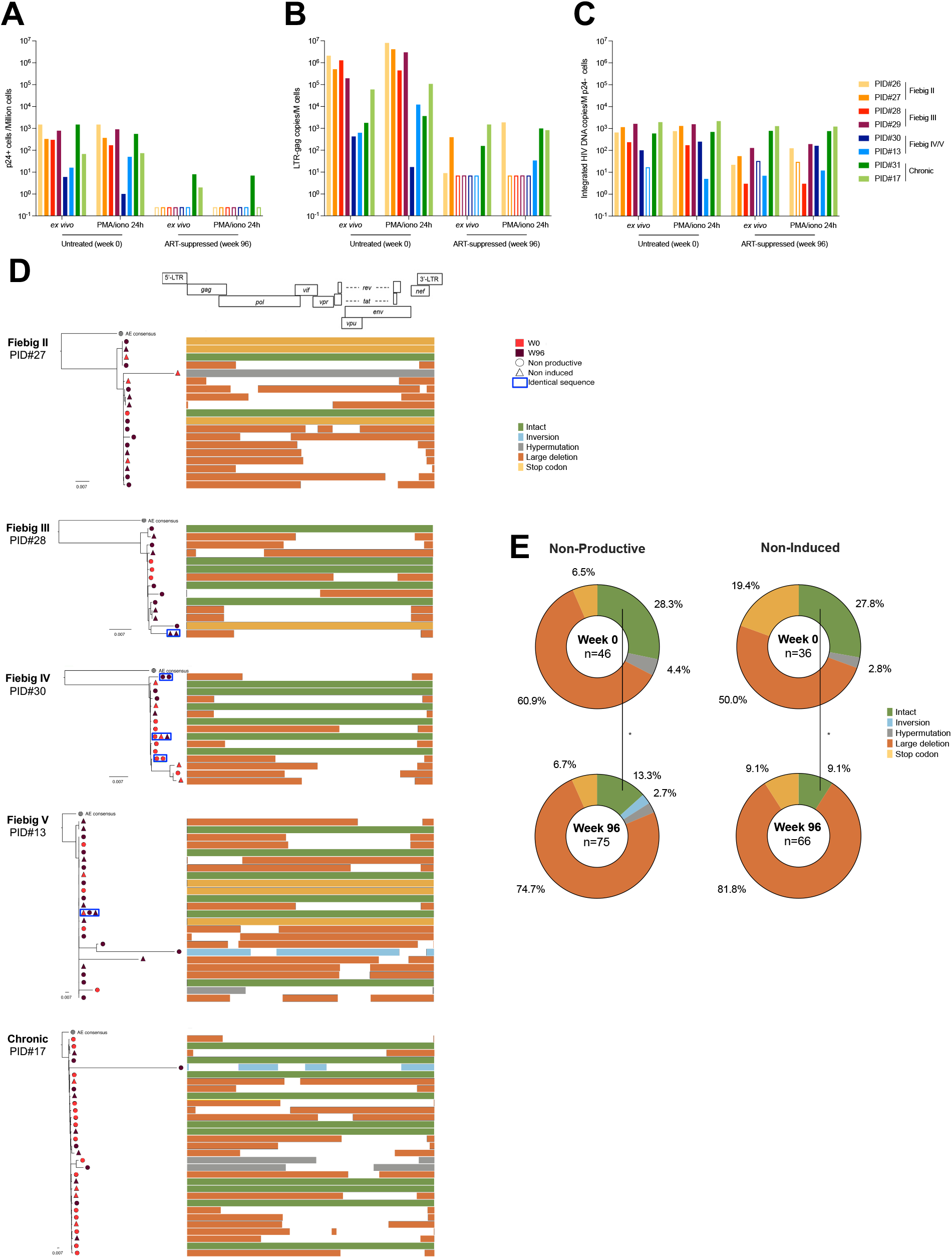
Inducibility, intactness and persistence of latent proviruses in acute infection and on ART. **A.** Frequency of p24+ cells measured by HIV-Flow in longitudinal blood samples of participants enrolled at different Fiebig stages (week 0) and after 96 weeks of ART. Frequencies were measured *ex vivo* to detect productively infected cells and after stimulation with PMA/ionomycin for 24h to detect the inducible reservoir. **B.** LTR-gag transcripts were measured in unstimulated or stimulated CD4+ T cells. RNA copies were normalized to the number of input cells. **C.** p24-cells were bulk sorted for integrated HIV DNA quantification by Alu-PCR. **A-C.** Empty bars represent undetectable measures, and the limit of detection is plotted. **D.** Near-full length genome amplification was performed in p24-cells to assess the intactness of latent genomes. To assess the inducibility of these proviruses, p24-cells from both the *ex vivo* and stimulation conditions were included. Phylogenetic trees representing the proviral landscape of five participants are depicted. Sequences obtained before (week 0, red) and during suppressive ART time point (week 96, burgundy) are included; Sequences obtained from p24-cells sorted directly *ex vivo* (circle) or after stimulation (triangle), representing the non-productive and the non-induced latent HIV reservoirs, respectively, are also included. Identical sequences are framed and represented on the same tree branch. On the right of the graph, proviral sequences are mapped to the HIV genome, and color-coded as follows: intact sequences in green, inversions in blue, hypermutations in grey, large deletions in orange and stop codons in yellow. **E.** Pie charts summarizing the frequency of defects and intact proviruses obtained from p24-cells sorted *ex vivo*(non-productive) and after stimulation (non-induced) before (week 0) and after ART initiation (week 96). The total number of sequences per condition is indicated at the center of the pie (sequence color categories are as in **D.**). Significant differences are highlighted (Wilcoxon or Fisher’exact test; p<0.05, *; p<0.01**; p<0.001, ***; p<0.0001 ****).

To determine if non-inducible proviruses could have been archived during acute infection, we bulk-sorted p24-cells at both time points and analyzed their proviral contents. Total and integrated HIV DNA were readily detected in almost all samples both during viremia and after 96 weeks of ART (Fig. 5c and Fig. S8e). Collectively, these data indicate that non-inducible proviruses are archived since the earliest stages of acute infection and can persist on ART. To assess the genetic integrity and persistence of these archived and non-induced proviruses, we performed near fulllength genome amplification in limiting dilutions of p24-cells lysates followed by PacBio sequencing. A total of 223 proviral sequences were obtained from the *ex vivo* and stimulated conditions, representing the non-productive (no spontaneous production of p24) and non-induced (no induction of p24 production upon stimulation) HIV-reservoirs, respectively. Intact noninduced proviruses were observed in 6/8 participants (PID #13, 17, 27-30; Fig. 5d and Fig. S8f) whereas only defective proviruses were detected in the last 2/8 (PID #26 and 31; Fig. S8f). During untreated infection, a third of the viral genomes retrieved from both non-induced and non-productively infected cells were genetically intact (Fig. 5e), indicating that deeply latent intact proviruses are archived since the earliest stage of infection. Although the proportion of noninduced intact genomes diminished after 2 years of ART (from 27.8% to 9.1%), they were still detected in most participants, demonstrating that early ART does not prevent the establishment of latent and genetically intact HIV genomes. Identical sequences were rare both before ART initiation and after 2 years of suppressive therapy (median, 0 and 5%, respectively; Fig. S8g), indicating that clonal expansions minimally contribute to the early dissemination and persistence of latent HIV genomes (p24-cells), similar to what we observed using TCR sequencing (Fig. 3a). Nonetheless, identical and genetically intact non-induced proviruses were found both in the pre- and post-ART samples from two participants (Participants #30 and #13, who initiated ART during Fiebig stages IV and V, respectively; Fig. 5d), indicating that genetically intact genomes that are refractory to viral reactivation are established early in infection and can persist during ART.

## Discussion

Using a combination of single-cell approaches, we extensively characterized the landscape of productively infected cells at the earliest stages of HIV infection. We observed that the frequency of these cells rapidly rises concomitant with plasma viremia and that their phenotype evolves throughout the different stages of acute infection. Using TCR and near-full length HIV genome sequencing, we demonstrate that multiple independent infection events both in blood and lymph nodes drive the production of a relatively homogeneous viral population during acute infection and that a latent pool of cells harboring intact HIV genomes is established early in infection and can persist during ART.

Using HIV-Flow, we measured the frequencies of productively infected cells in blood and lymph node during acute infection and observed that these frequencies rise from less than 10 to up to 1,000 per million CD4+ T cells (i.e. 100-fold) in a short interval (less than a week during the transition from Fiebig stage I to III(*26, 30*)), demonstrating the extremely fast dissemination of HIV and reflecting the kinetics of plasma viral load (*26*). This is in line with the results from a previous study conducted in individuals infected for a median time of 4 months in whom around 2,000 RNA+ cells/per million CD4+ T cells were quantified in lymph node (*31*). In our study, frequencies of p24+ cells in blood and lymph node were relatively similar, suggesting a possible recirculation of productively infected cells between these compartments (*32*), which is supported by the shared *env* sequences between the two sites.

When we analyzed the phenotype of productively infected cells, we observed that the vast majority of p24+ cells displayed a memory phenotype. Low cell surface expression levels of CD4 and MHC-I molecules suggest that *nef* was functional in most productively infected cells. The intriguing observation that not all p24+ cells expressed CCR5 suggests that this chemokine receptor may also be downregulated from the cell surface as previously proposed (*33*) or that HIV may use alternate coreceptors during acute infection (*34*). Unexpectedly, we observed that only 20-40% of p24+ cells expressed the proliferation marker Ki67, suggesting that HIV can productively infect cells that have not entered the cell cycle, as reported previously in several studies performed *in vitro* (*35, 36*).

The contribution of Tfh cells to the pool of productively infected cells was minimal throughout all stages of acute infection. While we cannot exclude that these cells were actually infected and immediately depleted, our observations argue against this model: We observed extremely low frequencies of Tfh in lymph nodes in acute infection, and their infection was concomitant with their development during chronic infection. Hence, our results are consistent with a model in which Tfh cells are scarce during acute infection, expand during early chronic infection when they become major targets for HIV replication. Therefore, our results suggest that Tfh cells have a limited role when plasma viremia peaks but are major contributors to the set point viremia. This is also supported by the delayed formation of germinal centers in SIV-infected macaques (*37*) and in HIV-infected individuals (*38*).

While we cannot exclude that changes in the phenotype of p24+ cells over time could be attributed to the modulation of the expression of cellular markers by HIV proteins, our data are consistent with a model in which HIV initially infects a small pool of proliferating cells expressing high levels of CCR5 in lymph nodes and then disseminates to other, possibly less permissive cell subsets. This is unlikely to be attributed to changes in the tropism of the viral populations since envelope sequences remain largely conserved throughout all stages of acute infection.

An unexpected finding was the fact that the overwhelming majority of productively infected cells expressed distinct TCRs during acute infection. This observation excludes two models: One in which an infected cell proliferates to disseminate HIV and one in which a clone proliferating actively at the time of infection supports viral dissemination. Instead, HIV targets individual cells belonging to distinct clonotypes that can be specific for several pathogens including HIV (*39*), *M. tuberculosis* (*40*) or influenzae (*41*). Since the repertoire of p24+ cells was heavily biased, it is likely that cells reactive to specific antigens and expressing specific TRBV and/or TRBJ regions were more susceptible or, on the contrary, more resistant to infection. For instance, in individuals from our cohort who had either been vaccinated or who have latent tuberculosis, we found that *M. tuberculosis*-specific cells were less frequently infected than cells from other specificities. This relative resistance to HIV infection may be attributed to high levels of MIP1-□ production by *M. tuberculosis*-specific cells in this particular context (*39, 40, 42*), as previously reported for CMV specific cells (*43*).

The fact that HIV preferentially infects cells belonging to large clones (*44*) that recirculate between blood and tissues is consistent with the preferential infection of effector memory CD4+ T cells during acute infection, as we previously reported (*8, 45*). Indeed, when compared to less differentiated central memory cells, effector memory cells are known to display restricted TCR repertoire as a result of their expansions (*46, 47*), to contain clones shared between anatomical sites (*48*), and to express higher levels of CCR5.

An outstanding question in the field of HIV reservoirs is how viral latency is established. While some studies have suggested that HIV latency can be directly established in resting CD4+ T cells (pre-activation latency) (*49, 50*), others have proposed that latency is primarily established when HIV infects cells that transition from an activated phenotype to a resting memory state (*20*). Although it is likely that both models coexist, our observations are consistent with a model in which latency can be directly established in a minority of the CD4+ T cells initially targeted by the virus: we found translationally inactive and genetically intact proviruses even at the earliest stage of infection (Fiebig II; before peak viremia). The vast majority of these latently infected cells resisted reactivation *in vitro*, suggesting that “deep latency” was already established at this early stage.

Within the pool of cells harboring non-inducible proviruses, those with intact proviruses decreased after 2 years of ART, indicating that they were preferentially cleared (*51*). This could suggest that a fraction of these intact proviruses underwent reactivation and were eliminated. Alternatively, intact non-induced genomes may have integrated into the chromatin of cells that are relatively short-lived compared to those in which defective proviruses were archived (*52*). As we observed in p24+ cells, duplication of HIV genomes was also rare in latently infected cells. We conclude that clonal expansion has a minimal contribution to the establishment of both the latently and productively infected pools of cells.

Our study has several limitations. First we did not analyze the early dissemination of HIV in the gut, which is thought to be very important for the establishment of productive infection and persistence on ART (*7, 53*). In addition, even after stimulation *in vitro*, we did not detect p24+ cells in samples from these early treated individuals after 2 years of ART: This precluded the study of the reactivatable reservoirs on ART. Although our data suggest that most of the latently infected cells persisting in these individuals carry proviruses that are difficult to reactivate, we previously observed that 8/8 Fiebig I participants experienced viral rebound upon treatment cessation (*15*). Our *in vitro* stimulation assay was not sensitive enough to recapitulate this rebound. Larger amounts of blood cells or access to tissue cells may be required to mimic viral reactivation. Alternatively, our *in vitro* stimulation may not have recapitulated the complex mechanisms that are causing viral resurgence *in vivo*.

In spite of these limitations, this study is, to our knowledge, the first report on the nature of the cells and the initial events that contribute to the dissemination of HIV in humans. We also report on the simultaneous establishment of two pools of productively and hard-to-reactivate latently infected cells. Therefore, our findings suggest that unlike pre- or post-exposure strategies which block early replication events, early ART will not prevent the establishment of CD4+ T cells harboring genetically intact and deeply latent proviruses.

## Supporting information

Auxilliary supplementary file

## Acknowledgements

The study team is grateful to the individuals who volunteered to participate in this study and the staff at SEARCH/IHRI, the Thai Red Cross AIDS Research Centre and the Department of Retrovirology, U.S. Army Medical Component, Armed Forces Research Institute of Medical Sciences (AFRIMS). We thank the RV254/SEARCH 010 and RV304/SEARCH Protocols team members. We thank the flow cytometry core at the CRCHUM, managed by Dominique Gauchat, Philippe St-Onge and Gaël Dulude for cell sorting, Corentin Richard for bioinformatics analysis as well as the NC3 core (Olfa Debbeche). We are grateful to the Thai Government Pharmaceutical Organization (GPO), ViiV Healthcare, Gilead Sciences and Merck for providing the antiretroviral medications for the RV254/SEARCH 010 study.

## Author contributions

Conceptualization: PG, AP, MP, CD, RF, JLM, LT, NC

Methodology: PG, AP, RF, NC

Investigation: SB, SP, EK, MLR, NP, JA, DH and SV

Visualization: PG

Funding acquisition: NC

Project administration: NC

Supervision: NC

Writing – original draft: PG, NC

Writing – review & editing: all authors

## Data and Materials Availability

All data generated or analyzed during this study are included in this published article (and its supplementary information files). Source data are provided with this paper within Supplementary Files. TCR sequences from CD4+ T cells and p24+ cells are provided as an Auxiliary Supplementary file. HIV *env* and HIV near full length sequences were submitted to GenBank. External databases used in this study are available online: IMGT® database (IMGT®, the international ImMunoGeneTics information system® [http://www.imgt.org]); McPAS-TCR database ([http://friedmanlab.weizmann.ac.il/McPAS-TCR/]).

## References

1. F. Barre-Sinoussi et al., Isolation of a T-lymphotropic retrovirus from a patient at risk for acquired immune deficiency syndrome (AIDS). Science 220, 868–871 (1983).

2. J. B. Whitney et al., Rapid seeding of the viral reservoir prior to SIV viraemia in rhesus monkeys. Nature 512, 74–77 (2014).

3. C. J. Miller et al., Propagation and dissemination of infection after vaginal transmission of simian immunodeficiency virus. J Virol 79, 9217–9227 (2005).

4. Z. Q. Zhang et al., Roles of substrate availability and infection of resting and activated CD4+ T cells in transmission and acute simian immunodeficiency virus infection. Proc Natl Acad Sci U S A 101, 5640–5645 (2004).

5. S. N. Gordon et al., Severe depletion of mucosal CD4+ T cells in AIDS-free simian immunodeficiency virus-infected sooty mangabeys. J Immunol 179, 3026–3034 (2007).

6. X. Wang et al., Massive infection and loss of CD4+ T cells occurs in the intestinal tract of neonatal rhesus macaques in acute SIV infection. Blood 109, 1174–1181 (2007).

7. J. D. Estes et al., Defining total-body AIDS-virus burden with implications for curative strategies. Nat Med 23, 1271–1276 (2017).

8. L. Leyre et al., Abundant HIV-infected cells in blood and tissues are rapidly cleared upon ART initiation during acute HIV infection. Sci Transl Med 12, (2020).

9. M. Perreau et al., Follicular helper T cells serve as the major CD4 T cell compartment for HIV-1 infection, replication, and production. J Exp Med 210, 143–156 (2013).

10. A. S. Perelson, A. U. Neumann, M. Markowitz, J. M. Leonard, D. D. Ho, HIV-1 dynamics in vivo: virion clearance rate, infected cell life-span, and viral generation time. Science 271, 1582–1586 (1996).

11. X. Wei et al., Viral dynamics in human immunodeficiency virus type 1 infection. Nature 373, 117–122 (1995).

12. M. L. Robb et al., Prospective Study of Acute HIV-1 Infection in Adults in East Africa and Thailand. N Engl J Med 374, 2120–2130 (2016).

13. V. H. Wu et al., Assessment of HIV-1 integration in tissues and subsets across infection stages. JCI Insight 5, (2020).

14. J. M. Coffin et al., Clones of infected cells arise early in HIV-infected individuals. JCI Insight 4, (2019).

15. D. J. Colby et al., Rapid HIV RNA rebound after antiretroviral treatment interruption in persons durably suppressed in Fiebig I acute HIV infection. Nat Med 24, 923–926 (2018).

16. P. Gantner et al., Single-cell TCR sequencing reveals phenotypically diverse clonally expanded cells harboring inducible HIV proviruses during ART. Nat Commun 11, 4089 (2020).

17. Z. Wang et al., Expanded cellular clones carrying replication-competent HIV-1 persist, wax, and wane. Proc Natl Acad Sci U S A 115, E2575–E2584 (2018).

18. P. Mendoza et al., Antigen-responsive CD4+ T cell clones contribute to the HIV-1 latent reservoir. J Exp Med 217, (2020).

19. F. R. Simonetti et al., Antigen-driven clonal selection shapes the persistence of HIV-1-infected CD4+ T cells in vivo. J Clin Invest 131, (2021).

20. L. Shan et al., Transcriptional Reprogramming during Effector-to-Memory Transition Renders CD4(+) T Cells Permissive for Latent HIV-1 Infection. Immunity 47, 766–775 e763 (2017).

21. S. Saleh et al., CCR7 ligands CCL19 and CCL21 increase permissiveness of resting memory CD4+ T cells to HIV-1 infection: a novel model of HIV-1 latency. Blood 110, 4161–4164 (2007).

22. P. U. Cameron et al., Establishment of HIV-1 latency in resting CD4+ T cells depends on chemokine-induced changes in the actin cytoskeleton. Proc Natl Acad Sci U S A 107, 16934–16939 (2010).

23. A. A. Okoye et al., Early antiretroviral therapy limits SIV reservoir establishment to delay or prevent post-treatment viral rebound. Nat Med 24, 1430–1440 (2018).

24. M. S. De Souza et al., Impact of nucleic acid testing relative to antigen/antibody combination immunoassay on the detection of acute HIV infection. AIDS 29, 793–800 (2015).

25. M. Pardons et al., Single-cell characterization and quantification of translation-competent viral reservoirs in treated and untreated HIV infection. PLoS Pathog 15, e1007619 (2019).

26. E. W. Fiebig et al., Dynamics of HIV viremia and antibody seroconversion in plasma donors: implications for diagnosis and staging of primary HIV infection. AIDS 17, 1871–1879 (2003).

27. J. Lama, A. Mangasarian, D. Trono, Cell-surface expression of CD4 reduces HIV-1 infectivity by blocking Env incorporation in a Nef- and Vpu-inhibitable manner. Curr Biol 9, 622–631 (1999).

28. P. Meysman et al., On the viability of unsupervised T-cell receptor sequence clustering for epitope preference. Bioinformatics 35, 1461–1468 (2019).

29. D. A. Bolotin et al., MiXCR: software for comprehensive adaptive immunity profiling. Nat Methods 12, 380–381 (2015).

30. J. Ananworanich et al., A novel acute HIV infection staging system based on 4th generation immunoassay. Retrovirology 10, 56 (2013).

31. T. Schacker et al., Productive infection of T cells in lymphoid tissues during primary and early human immunodeficiency virus infection. J Infect Dis 183, 555–562 (2001).

32. T. T. Murooka et al., HIV-infected T cells are migratory vehicles for viral dissemination. Nature 490, 283–287 (2012).

33. M. Toyoda et al., Differential Ability of Primary HIV-1 Nef Isolates To Downregulate HIV-1 Entry Receptors. J Virol 89, 9639–9652 (2015).

34. J. Isaacman-Beck et al., Heterosexual transmission of human immunodeficiency virus type 1 subtype C: Macrophage tropism, alternative coreceptor use, and the molecular anatomy of CCR5 utilization. J Virol 83, 8208–8220 (2009).

35. A. Shen et al., Endothelial cell stimulation overcomes restriction and promotes productive and latent HIV-1 infection of resting CD4+ T cells. J Virol 87, 9768–9779 (2013).

36. L. M. Agosto, M. B. Herring, W. Mothes, A. J. Henderson, HIV-1-Infected CD4+ T Cells Facilitate Latent Infection of Resting CD4+ T Cells through Cell-Cell Contact. Cell Rep 24, 2088–2100 (2018).

37. J. J. Hong, P. K. Amancha, K. Rogers, A. A. Ansari, F. Villinger, Spatial alterations between CD4(+) T follicular helper, B, and CD8(+) T cells during simian immunodeficiency virus infection: T/B cell homeostasis, activation, and potential mechanism for viral escape. J Immunol 188, 3247–3256 (2012).

38. J. L. Mitchell et al., Anti-HIV antibody development up to 1 year after antiretroviral therapy initiation in acute HIV infection. J Clin Invest 132, (2022).

39. D. C. Douek et al., HIV preferentially infects HIV-specific CD4+ T cells. Nature 417, 95–98 (2002).

40. C. Geldmacher et al., Preferential infection and depletion of Mycobacterium tuberculosis-specific CD4 T cells after HIV-1 infection. J Exp Med 207, 2869–2881 (2010).

41. R. B. Jones, C. Kovacs, T. W. Chun, M. A. Ostrowski, Short communication: HIV type 1 accumulates in influenza-specific T cells in subjects receiving seasonal vaccination in the context of effective antiretroviral therapy. AIDS Res Hum Retroviruses 28, 1687–1692 (2012).

42. T. Dragic et al., HIV-1 entry into CD4+ cells is mediated by the chemokine receptor CC-CKR-5. Nature 381, 667–673 (1996).

43. J. P. Casazza et al., Autocrine production of beta-chemokines protects CMV-Specific CD4 T cells from HIV infection. PLoS Pathog 5, e1000646 (2009).

44. J. A. Collora et al., Single-cell multiomics reveals persistence of HIV-1 in expanded cytotoxic T cell clones. Immunity, (2022).

45. J. Grau-Exposito et al., A Novel Single-Cell FISH-Flow Assay Identifies Effector Memory CD4(+) T cells as a Major Niche for HIV-1 Transcription in HIV-Infected Patients. mBio 8, (2017).

46. I. Bretschneider et al., Discrimination of T-cell subsets and T-cell receptor repertoire distribution. Immunol Res 58, 20–27 (2014).

47. E. A. Boritz et al., Multiple Origins of Virus Persistence during Natural Control of HIV Infection. Cell 166, 1004–1015 (2016).

48. M. Miron et al., Maintenance of the human memory T cell repertoire by subset and tissue site. Genome Med 13, 100 (2021).

49. V. A. Evans et al., Myeloid dendritic cells induce HIV-1 latency in non-proliferating CD4+ T cells. PLoS Pathog 9, e1003799 (2013).

50. L. Chavez, V. Calvanese, E. Verdin, HIV Latency Is Established Directly and Early in Both Resting and Activated Primary CD4 T Cells. PLoS Pathog 11, e1004955 (2015).

51. M. R. Pinzone et al., Longitudinal HIV sequencing reveals reservoir expression leading to decay which is obscured by clonal expansion. Nat Commun 10, 728 (2019).

52. J. Neidleman et al., Phenotypic analysis of the unstimulated in vivo HIV CD4 T cell reservoir. Elife 9, (2020).

53. Q. Li et al., Peak SIV replication in resting memory CD4+ T cells depletes gut lamina propria CD4+ T cells. Nature 434, 1148–1152 (2005).

54. M. P. Lefranc et al., IMGT, the international ImMunoGeneTics database. Nucleic Acids Res 27, 209–212 (1999).

55. N. Tickotsky, T. Sagiv, J. Prilusky, E. Shifrut, N. Friedman, McPAS-TCR: a manually curated catalogue of pathology-associated T cell receptor sequences. Bioinformatics 33, 2924–2929 (2017).

56. B. Hiener et al., Identification of Genetically Intact HIV-1 Proviruses in Specific CD4(+) T Cells from Effectively Treated Participants. Cell Rep 21, 813–822 (2017).

