## Supplementary, Figures and Tables for "HIV rapidly targets a diverse pool of CD4+ T cells to establish productive and latent infections"

**Gantner et al.**

### Methods

##### Participants and sample collection

The RV254/SEARCH 010 cohort study (clinicaltrials.gov NCT00796146) enrolls participants at the earliest stages of acute HIV infection at the Thai Red Cross AIDS Research Centre in Bangkok. High-risk volunteers are screened for acute HIV infection (AHI) in real time with pooled nucleic acid testing and sequential immunoassay (IA). Individuals with AHI were enrolled if they had a positive HIV RNA with or without a reactive IA. Participants were then categorized as Fiebig stages I to V as follows: Fiebig I - positive HIV RNA, negative p24 antigen, non-reactive 3$^{rd}$ generation IA; Fiebig II – positive HIV RNA, positive p24 antigen, non-reactive 3$^{rd}$ generation IA; Fiebig III - positive HIV RNA, positive p24 antigen, reactive 3$^{rd}$ generation IA, negative western blot; Fiebig IV - positive HIV RNA, positive or negative p24 antigen, reactive 3$^{rd}$ generation IA, indeterminate western blot; Fiebig V - positive HIV RNA, positive or negative p24 antigen, reactive 3$^{rd}$ generation IA, positive western blot except p31(*26*). ART was voluntary and offered to all participants and was initiated at a median (IQR) of 2 (0-5) days after enrollment (*8*) under a separate protocol (NCT00796263). Samples from chronically infected naïve individuals enrolled in the RV304/SEARCH 013 (*8*) (NCT01397669) studies were also obtained.

For the present study, paired PBMCs and LNMCs samples from 25 untreated individuals were analyzed. Eight participants also provided before and during suppressive ART (96 weeks) PBMCs samples. Both PBMCs and LNMCs were isolated by Ficoll density gradient centrifugation and were cryopreserved in liquid nitrogen.

##### Ethics statement

All clinical studies were approved by the Institutional Review Boards (IRBs) of Chulalongkorn University in Thailand, Walter Reed Army Institute of Research (WRAIR)
and the Centre Hospitalier de l'Université de Montréal in Canada. All participants gave written informed consent.

##### Cell culture

The HIV-Flow assay was used to quantify and analyze the phenotype of cells expressing p24 protein (*25*). Briefly, CD4+ T cells were isolated by negative magnetic selection using the EasySep Human CD4+ T Cell Enrichment Kit (StemCell Technology, Cat#19052). Purity was typically >98%. 5-15x$10^6$ CD4+ T cells were resuspended at 2x$10^6$ cells/mL in RPMI + 10% Fetal Bovine Serum and antiretroviral drugs were added to the culture medium (200nM raltegravir, 200nM lamivudine). Cells were then rested (18h) or stimulated, depending on the experiment. Stimulation included a pre-incubation of 1h with 5µg/mL Brefeldin A (BFA, Sigma, Cat#B2651) before stimulation in order to prevent the upregulation of cell surface markers, and BFA was maintained in the culture until the end of the stimulation. Cells were then stimulated with 1µg/mL ionomycin (Sigma, Cat#I9657) and 162nM PMA (24h) (Sigma, Cat#P8139).

**Flow cytomety**

After resting or stimulation, cells were collected, resuspended in PBS and stained with the Aqua Live/Dead staining kit for 20min at 4°C. Cells were then stained with antibodies against extracellular molecules in PBS + 4% human serum (Atlanta Biologicals, Cat#540110) for 20min at 4°C. After a 45min fixation/permeabilization step was performed with the FoxP3 Transcription Factor Staining Buffer Set (eBioscience, Cat#00-5523-00) following the manufacturer's instructions, cells were then stained with anti-p24 KC57 and anti-p24 28B7 antibodies for an additional 45min at room temperature in the FoxP3 Buffer. Cells were then washed and resuspended in PBS for subsequent cell sorting.

p24 KC57-PE was purchased from Beckman Coulter (Cat#6604667, Dilution 1/1000) and p24 28B7-APC was purchased from MediMabs (Cat#MM-0289-APC, Dilution 1/1000). CD4-APC-H7 (Clone: RPA-T4, Cat#560168, Dilution 1/100), CD45RA-A700 (Clone: HI100, Cat#560673, Dilution 1/50), CXCR5-BB515 (Clone: RF8B2, Cat#564624, Dilution 1/50), CXCR3-PerCP-Cy5.5 (Clone: 1C6/CXCR3, Cat#560832, Dilution 1/25), Ki67-PE-Cy7 (Clone: B56, Cat#561283, Dilution 1/50), CCR5-BV421 (Clone: 2D7/CCR5 Cat#562576, Dilution 1/25), CD45RA-BV786 (Clone: HI100, Cat#563870, Dilution 1/50), HLA-DR-BV605 (Clone: G46-6, Cat#562845, Dilution 1/50), CCR7-BB515 (Clone: 3D12, Cat#565870, Dilution 1/25), were purchased from BD Bioscience. PD-1-BV605 (Clone: EH12.2H7, Cat#329923, Dilution 1/25), ICOS-BV785 (Clone: C398.4A, Cat#313534, Dilution 1/25), HLA-ABC-A700 (Clone: W6/32, Cat#311437, Dilution

1/100), CD69-PerCP-Cy5.5 (Clone: FN50, Cat#310926, Dilution 1/50), Bcl-2-BV421 (Clone: 100, Cat#658709, Dilution 1/50), were purchased from BioLegend. Live/Dead Aqua Cell Stain (405nm) was purchased from ThermoFisher Scientific (Cat#L34957).

**Flow cytometry cell sorting**

The frequency of p24 double positive cells (KC57+, 28B7+) was determined by flow cytometry in gated viable T cells. Examples of gating strategies are represented in Fig. S1 and S9. In all experiments, CD4+ T cells from an HIV-uninfected control were included to set the threshold of positivity. Single p24 double positive (p24+ cells) were indexed-sorted and p24 double negative (p24- cells) were bulk sorted on a BD FACS ARIA III. Single-cells were sorted in 96-wells PCR plates containing 7.6µL of DirectPCR Lysis Reagent (Viagen Biotech) and 0.4 µL of 10mg/mL proteinase K (from Wisent, 25530–015). The PCR plates were subsequently incubated at 55°C for 1 hour for cell lysis followed by 10 min at 95°C to inactivate proteinase K. Bulk-sorted and pelleted cells were digested in 30µL of Proteinase K (Invitrogen#25530-015, final concentration 400 µg/ml, TriS HCl (10mM) and KCl (50nM) overnight at 55°C followed by 10 min at 95°C to inactivate proteinase K. Flow cytometry data of p24+ cells were analyzed using FlowJo version 10.7.1. Cell lysates were then used for TCR sequencing or near-full length HIV genome sequencing.

**TCR amplification on genomic DNA**

We developed a two-step PCR method to amplify a portion of approximately 260bp of the TCRβ encompassing: (1) the end of the V segment, (2) the CDR3, and (3) the J segment, on genomic DNA from lysed single-cells. We used a set of 22 forward primers complementary to the 23 functional V segments families, and 13 reverse primers complementary to the 13 functional J segments, to amplify the target portion of the TCRβ in a first multiplex PCR reaction, as previously described (*16*). M13 forward and reverse tags were added to the 5' end of these primers, to allow a second PCR amplification, which was followed by Sanger sequencing. Sequences of all primers are listed in Table S2. The first PCR reaction was performed using the Qiagen Multiplex PCR kit (Cat#206143, Qiagen), in a total volume of 50 µL: 25 µL of Qiagen Multiplex PCR master mix, 10 µL of a mix of all primers (each primer at a concentration of 1.25 µM in the mix, providing a final concentration of 250 nM per primer), 5 µL of Q-Solution, and 10 µL of the single-cell lysate. First PCR conditions were as follows: 15 min at 95°C followed by 40 cycles of; 30 s at 95°C, 90 s at 68°C and 20 s at 72°C; with a final elongation for 5 min at 72°C. A second round of PCR reaction was performed using the M13F and M13R primers (see Table S2) and the Taq DNA Polymerase kit (Invitrogen), in a total volume of 50 μL: 5 μL of 10X PCR buffer, 3 μL of $MgCl_2$ (50 mM), 1.5 μL of dNTPs (10 mM), 2 μL of M13F primer and 2 μL of M13R primer (each at 10 μM, providing a final concentration of 400 nM per primer), 0.5 μl Taq DNA Polymerase (5 U/ μL), 26 μL $H_2O$ and 10 μL of the first PCR products. The amplification conditions for the second PCR reaction were as follows: 3 min at 94°C followed by 40 cycles of; 45 s at 94°C, 60 s at 55°C and 30 s at 72°C; with a final elongation of 10 min at 72°C. A third round PCR adding next-generation sequencing adaptors to the TCR amplicons was performed for bulk cells to allow for further MiSeq sequencing. The third PCRs were performed using the same amplification conditions as for the second one (see MiSeq adaptors primers in Table S2).

**TCR sequencing and analysis**

Successful amplification of the TCRβ region was verified by electrophoresis on a 2% agarose gel and followed by gel purification of the TCRβ bands using the Buffer QG and the QIAquick 96 PCR Purification kit (Cat#28181, Qiagen), according to the manufacturer's instructions. Sanger sequencing was performed by Eurofins Genomics, with M13F and M13R as sequencing primers. TCRβ sequences were re-constructed using both forward and reverse sequences, and were analyzed using the V-QUEST tool of the IMGT® database (IMGT®, the international ImMunoGeneTics information system® [http://www.imgt.org](*54*)) to retrieve TCRβ information, including V and J segments usage and junction/CDR3 analysis. Next-generation sequencing (MiSeq, Illumina) was performed by Genome Québec. TCR reads were aligned, identified and grouped using the regular pipeline of MiXCR version 3.0.13 (*29*). Clonotypes with less than 10 reads were removed, and clonotypes representing at least 0.1% of the total read count for each sample were considered as expanded. TCR sequences were analyzed using an algorithm to predict antigen specificity: CDR3 sequences were compared to the McPAS-TCR database of TCRs of known antigenic specificity ([http://friedmanlab.weizmann.ac.il/McPAS-TCR/](*55*)) and sequence similarities were identified. We predicted TCR specificity using the three criteria described by Meysman et al. (*28*): 1) CDR3 sequences should have identical length, 2) CDR3 sequences should be long enough and 3) CDR3 sequences should not differ by more than one amino acid. Among all CDR3 sequences, those fulfilling these three criteria with matched CDR3 sequences from the database were considered at high probability of sharing the same specificity.

**Near-full length HIV genome amplification on genomic DNA**

We developed a two-step PCR method to amplify a portion of approximately 9,000bp of the HIV genome on lysed bulk cells. Sequences of primers are listed in Table S2. Both PCR reactions were performed using the Platinum SuperFi II Master Mix (Cat#12368010, Life Tech). The first PCR was prepared in a total volume of 40 μL: 20 μL of Platinum SuperFi II master mix, 0.8 μL of each primer (each primer at a concentration of 10 μM in the mix, providing a final concentration of 200 nM per primer), 8.4 μL $H_2O$, and 10 μL of the cell lysate. First PCR conditions were as follows: 30 s at 98°C followed by 25 cycles of; 30 s at 98°C, 10 s at 60°C and 5 min at 72°C; with a final elongation for 5 min at 72°C. A nested PCR, with PacBio barcoded primers was then performed prior to PacBio sequencing. The nested PCR was prepared in a total volume of 30 μL: 15 μL of Platinum SuperFi II master mix, 0.6 μL of each primer (each primer at a concentration of 10 μM in the mix, providing a final concentration of 200 nM per primer), 8.8 μL $H_2O$, and 5 μL of the first PCR products diluted 1/3. Nested PCR conditions were as follows: 30 s at 98°C followed by 30 cycles of; 30 s at 98°C, 10 s at 60°C and 5 min at 72°C; with a final elongation for 5 min at 72°C.

**Near-full length HIV genome sequencing and analysis**

Successful amplification of the HIV genomes was verified by electrophoresis on a 0.5% agarose gel, followed by AMPure XP (Cat#A63881, Beckman Coulter) purification with a beads/PCR products ratio of 1.6. Purified amplicons were then quantified using Nanodrop (ThermoFischer) and pooled. PacBio sequencing was performed by Genome Québec on a PacBio Sequel II instrument. Obtained sequences were then demultiplexed and those blasting the HIV genome, with at least 30 reads, containing both primers sequences on their ends and clustering together within the same individual were then aligned. Alignments were then used for phylogeny and clonality analysis using Maximum-Likelihood tree GTR+I+G model, with 1000 bootstraps in iqtree2, version 2.1.2. Trees were annotated with FigTree version 1.4.4. Sequences were also analyzed for integrity using HIVDatabase QCTool (https://www.hiv.lanl.gov/content/sequence/QC/index.html) and the ProseqIT (https://psd.cancer.gov/tools/pvs_annot.php). Start and stop codons were confirmed using GeneCutter tool (https://www.hiv.lanl.gov/content/sequence/GENE_CUTTER/cutter.html), and the integrity of the packaging signal was manually determined. Defects were evaluated in the following order: inversion, hypermutation, length (large deletion, i.e. <8800 pb), stop codon, frameshift, packaging signal (ψ), as previously described (*56*). If no defects were found (excluding defects in *nef* and in *tat2*), the sequence was considered genetically intact.

**Data representations and statistical analyses**

Data were analyzed and represented using Graphpad Prism version 9.1.0. Results were represented as median or mean values, with interquartile range or minimum and maximum values, as indicated in the figure legends. Correlations were determined using nonparametric Spearman's test. For group comparisons, non-parametric Wilcoxon matched-pairs signed rank tests were used. P values of less or equal to 0.05 were considered statistically significant.

**Fig. S1**

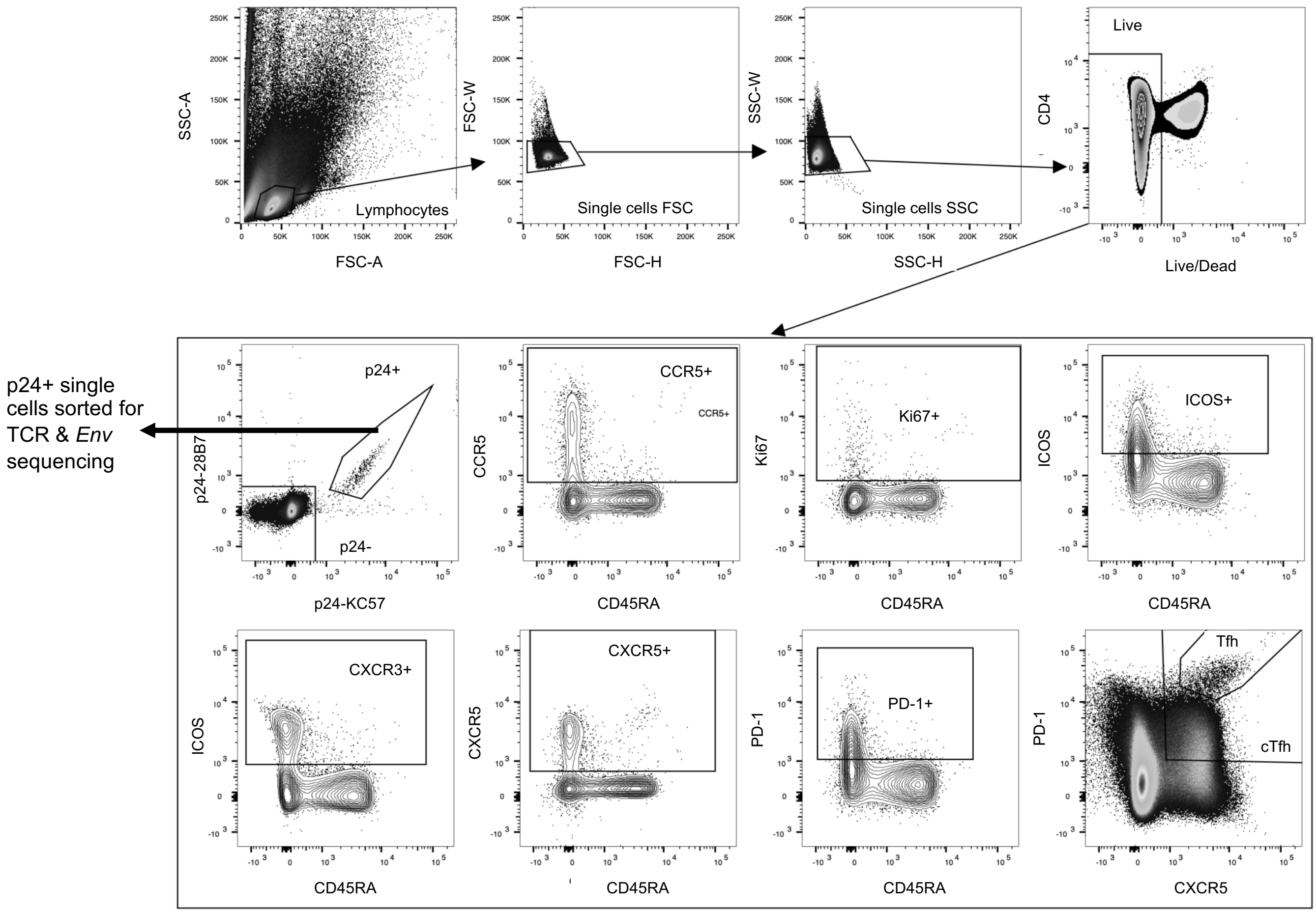


**Fig. S1. Gating strategy for single cell sorting.** Gating strategy used for index cell sorting of p24+ cells for subsequent TCR and HIV *Env* sequencing with recording of CD45RA, CCR5, Ki67, ICOS, CXCR3, CXCR5 and PD-1 expression levels.

Fig. S2

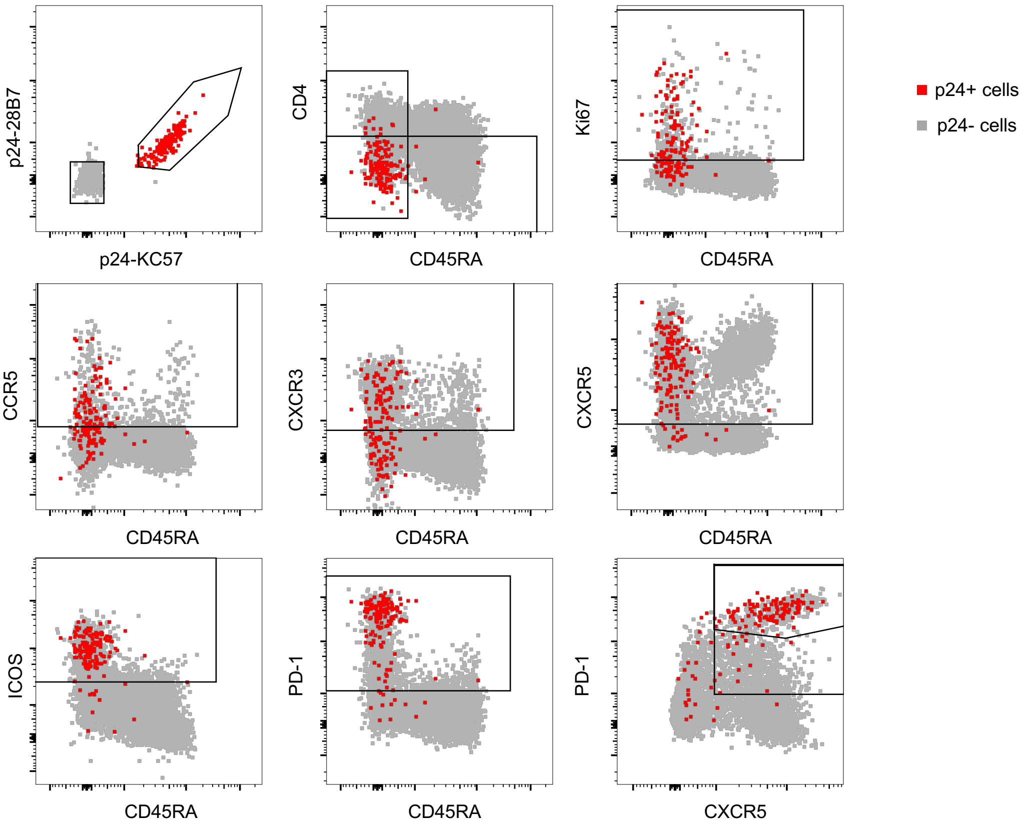


**Fig. S2. Phenotyping of p24+ cells.** Representative dot plots showing expression levels of CD4, CD45RA, Ki67, CCR5, CXCR3, CXCR5, ICOS and PD-1 in p24+ cells (in red), overlaid onto p24- cells (in grey).

**Fig. S3**

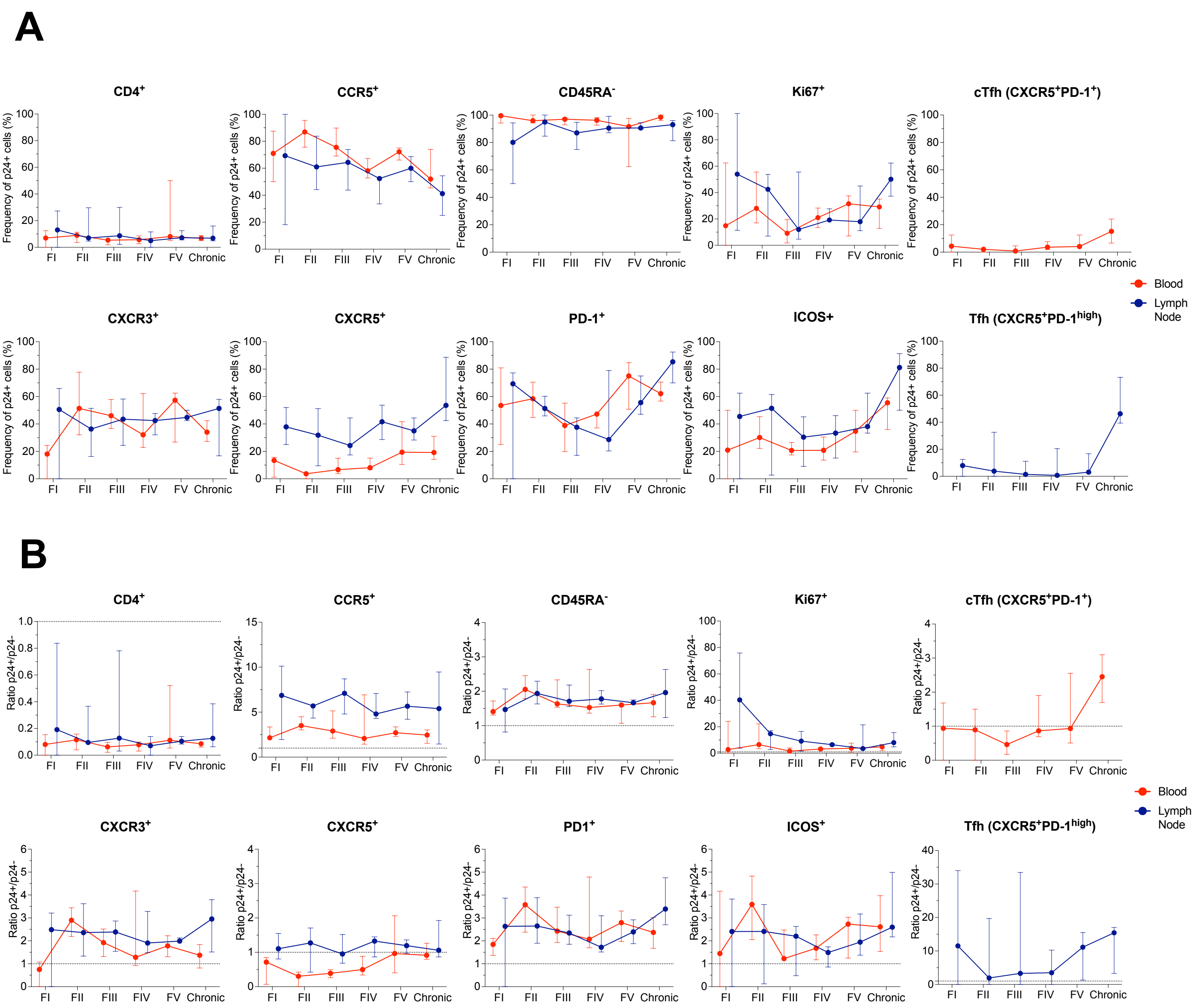


**Fig. S3. Expression levels of several cellular markers on p24+ cells at different Fiebig stages. A.** The frequency of p24+ or **B.** ratio of frequency (p24+/p24-) cells from blood (red) or lymph node (blue) expressing each marker or combination of markers (CD4+, CCR5+, CD45RA-, Ki67+, CXCR3+, CXCR5+, PD-1+, ICOS+, circulating T follicular helpers (cTfh) cells and Tfh cells) is depicted for each Fiebig stage (I to V and Chronic infection). Median values are plotted with 95% CI.

### Fig. S4

**A**

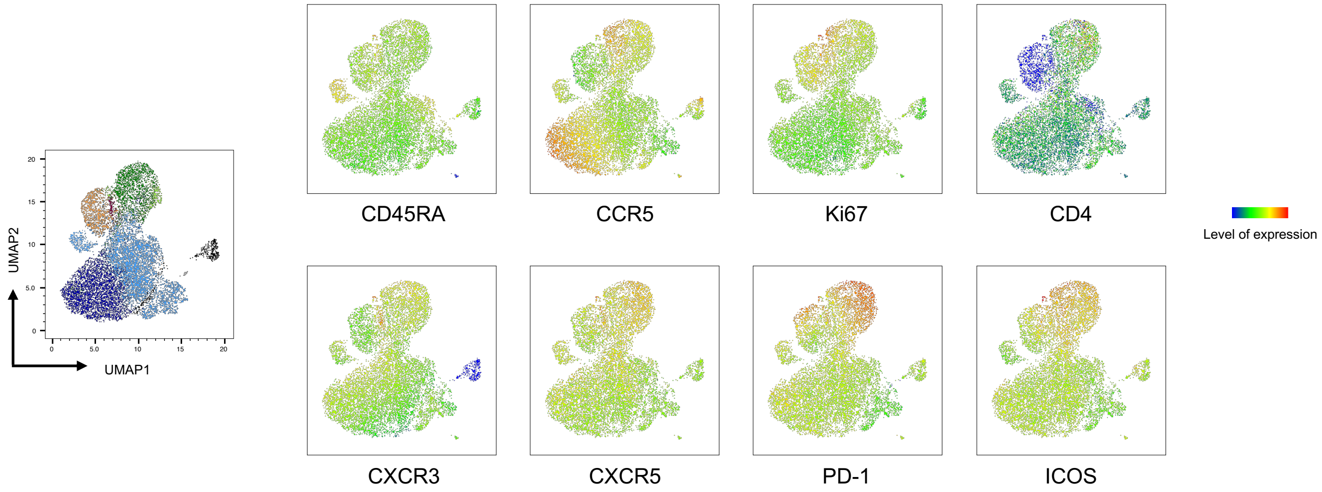


| | Cluster | Ki67 | CCR5 | CXCR3 | CXCR5 | PD-1 | ICOS |
|---|---|---|---|---|---|---|---|
| 1 | Tfh | +/- | ++ | +/- | ++ | +++ | + |
| 2 | Proliferative Tfh | +++ | +++ | +/- | + | ++ | ++ |
| 3 | Proliferative, Th1-like | +++ | + | +++ | ++ | +++ | ++ |
| 4 | Proliferative non-Th1-like | + | + | - | +/- | +/- | +/- |
| 5 | Non-Proliferative, Th1-like | - | +++ | + | +/- | + | + |
| 6 | Non-Proliferative non-Th1-like | - | +/- | - | +/- | +/- | +/- |
| 7 | Non-Th1 | - | + | - | - | + | + |
| 8 | Others | - | +/- | - | - | +/- | +/- |

**B**

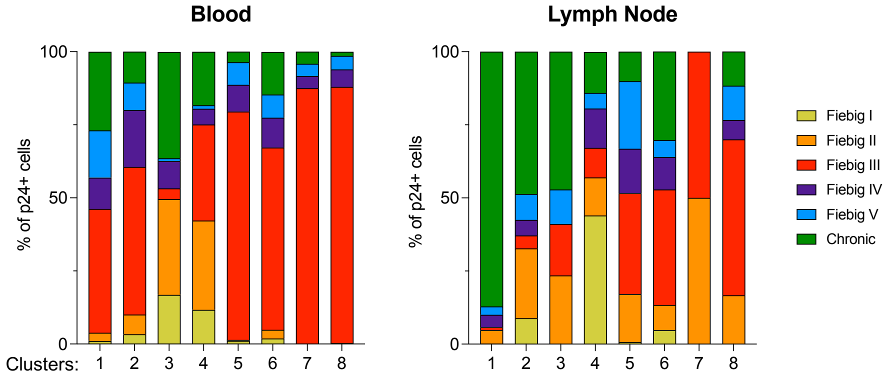


**Fig. S4. Expression of cellular markers in p24+ cells clusters. A.** p24+ cells phenotypic data were integrated in a UMAP analysis and generated 8 cell clusters (left). Relative levels of expression of CD45RA, CCR5, Ki67, CD4, CXCR3, CXCR5, PD-1 and ICOS is depicted. The table shows the phenotype of each cell cluster. **B.** Fiebig stages representation frequency among the 8 p24+ cell clusters depicted in bar graphs for both blood and lymph node.

#### Fig. S5

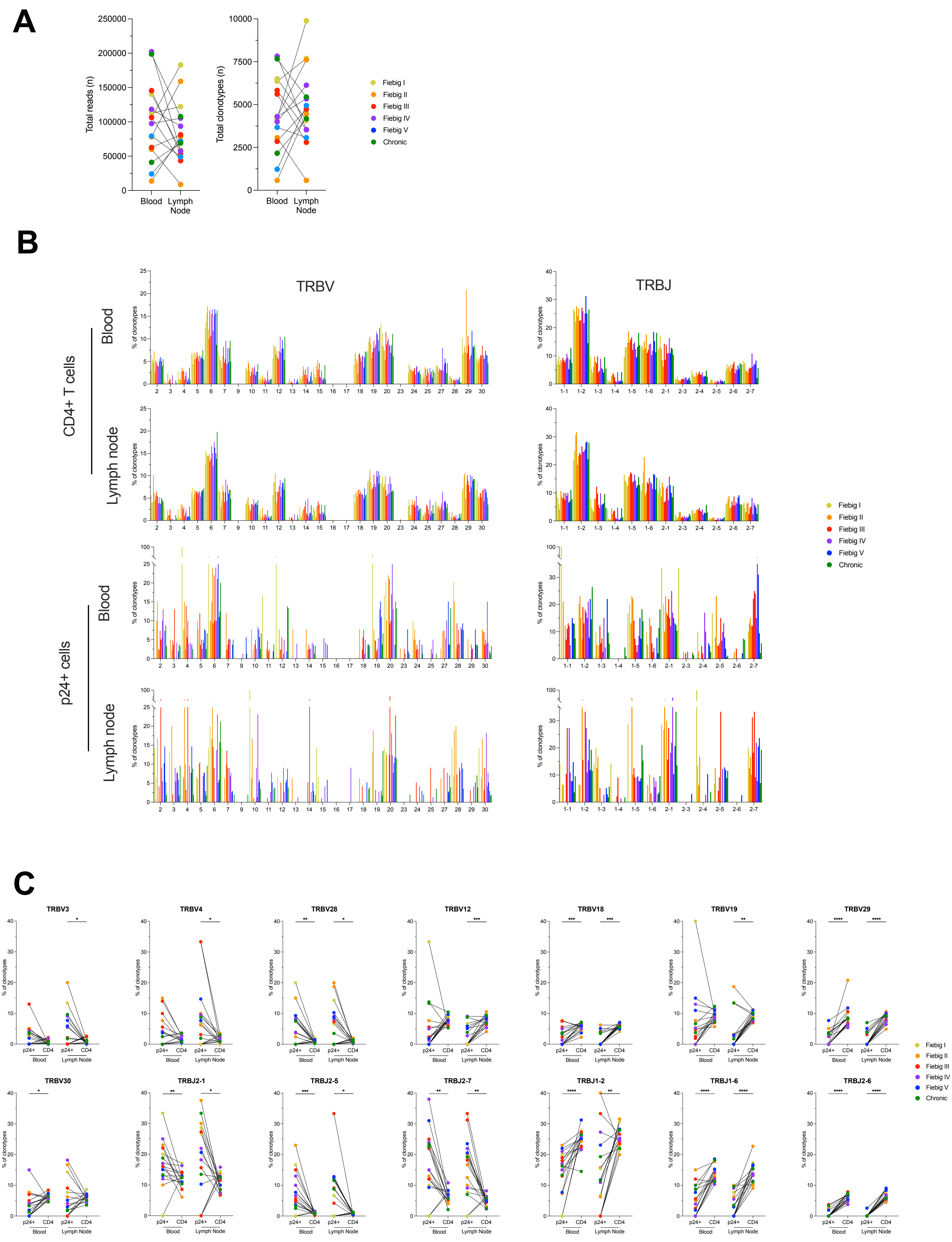
A
Total reads (n)
Total clonotypes (n)
Blood
Lymph Node
Fiebig I
Fiebig II
Fiebig III
Fiebig IV
Fiebig V
Chronic
B
TRBV
TRBJ
CD4+ T cells
p24+ cells
Blood
Lymph node
% of clonotypes
C
TRBV3
TRBV4
TRBV28
TRBV12
TRBV18
TRBV19
TRBV29
TRBV30
TRBJ2-1
TRBJ2-5
TRBJ2-7
TRBJ1-2
TRBJ1-6
TRBJ2-6
p24+
CD4
Blood
Lymph Node

**Fig. S5. Comparisons of TCR repertoire in blood and lymph nodes and in p24+ and total CD4+ T cells. A.** Number of TCR reads and clonotypes in paired blood and lymph nodes samples retrieved after TCR bulk sequencing of total CD4+ T cells. **B.** Frequency of TRBV and TRBJ segment usage for the clonotypes identified by TCRβ sequencing in p24+ cells and in total CD4+ T cells in both blood and lymph nodes. Each bar represents a single participant and is colored according to the stage of infection. **C.** Significant differences in the frequencies of TRBV and TRBJ segment usage for the clonotypes identified by TCRβ sequencing in p24+ cells and in total CD4+ T cells. (Wilcoxon; $p<0.05$, *; $p<0.01$**; $p<0.001$, ***; $p<0.0001$ ****).

**Fig. S6**

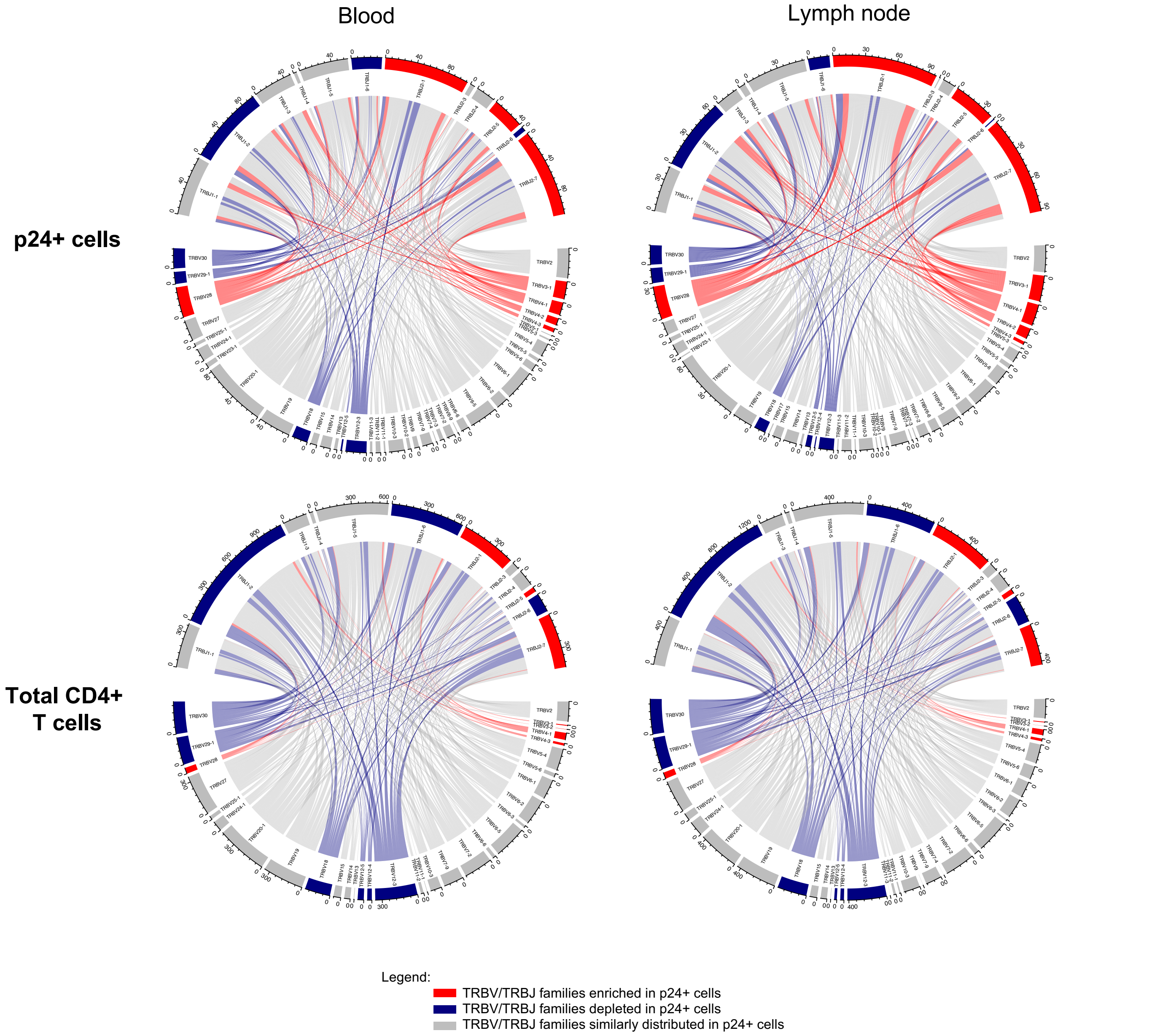


**Fig. S6. Association between the TRBV and TRBJ segments in p24+ and total CD4+ cells from blood and lymph nodes from all participants.** The circular axis represents the number of clonotypes in each TRBV (bottom) and TRBJ gene families (top). TRBV and TRBJ genes that were significantly increased/decreased in p24+ cells are highlighted with a red or blue background, respectively and so is the association link starting from the TRBV gene.

Fig. S7

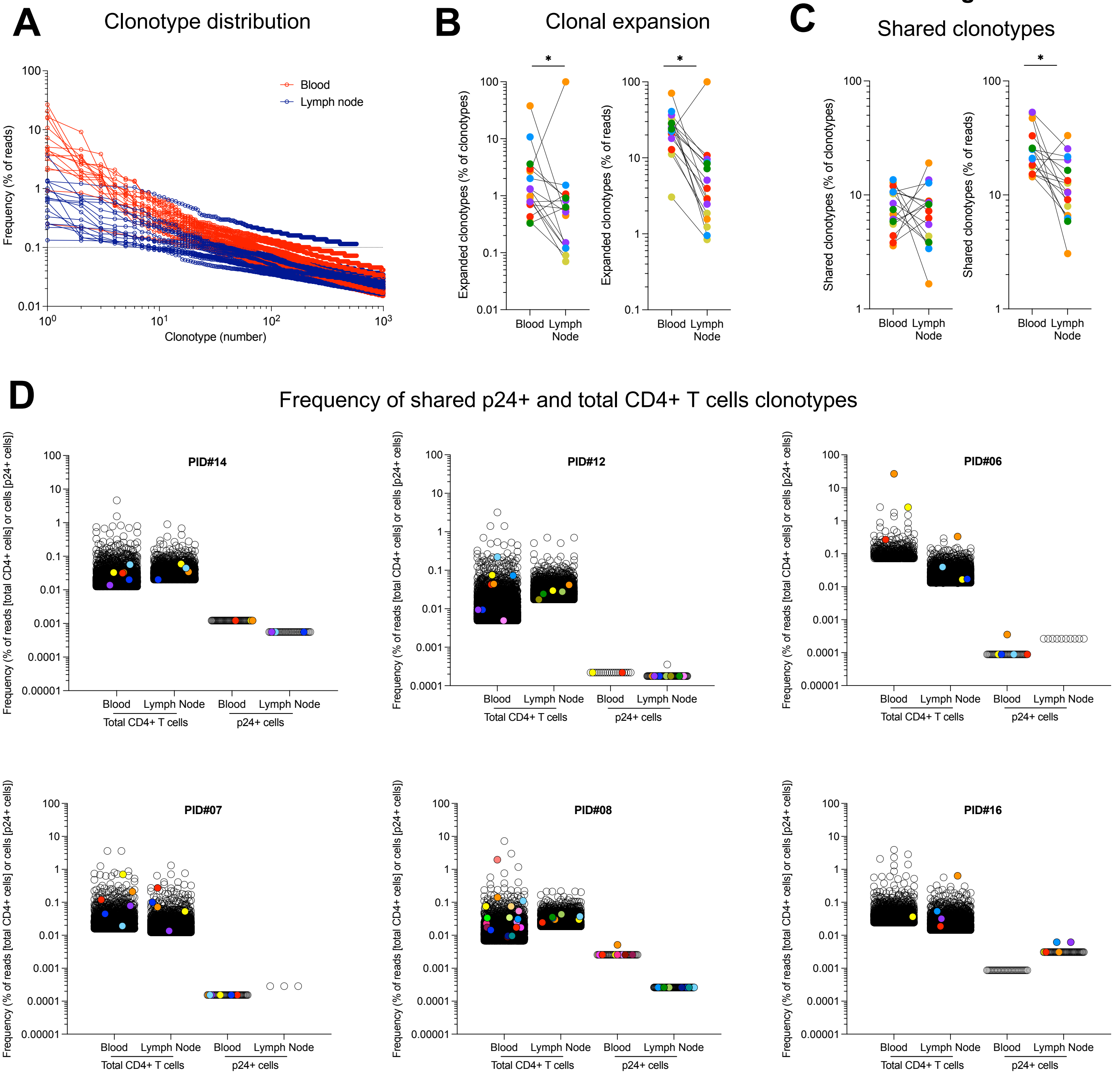


**Fig. S7. Distribution of p24+ and total CD4+ T cells TCR repertoires in blood and lymph nodes. A.** Distribution of clonotypes in total CD4+ T cells from blood and lymph nodes in all participants. Frequencies are shown as percentage of total reads. Dotted line represents the 0.1% frequency threshold. **B.** Frequency of the number and of the percentage of reads of likely expanded clonotypes (>0.1% of total reads) among all detected clonotypes in paired blood and lymph nodes samples, respectively. **C.** Common clonotypes between blood and lymph node samples. Frequency of  clonotypes and of reads of common blood/lymph node clonotypes among all clonotypes detected, in blood and lymph node samples, respectively. **D.** Distribution (based on bulk deep sequencing data and single-cell sorting/Sanger sequencing) of clonotypes corresponding to the clones that were found in a single subset (empty circles) and in multiple subsets including p24+ cells (colored circles). Frequencies are shown as percentage of total reads for total CD4+ T cells and as the frequency of cells determined by HIV-Flow for p24+ cells. (Wilcoxon; $p<0.05$, *; $p<0.01$**; $p<0.001$, ***; $p<0.0001$ ****).

Fig. S8

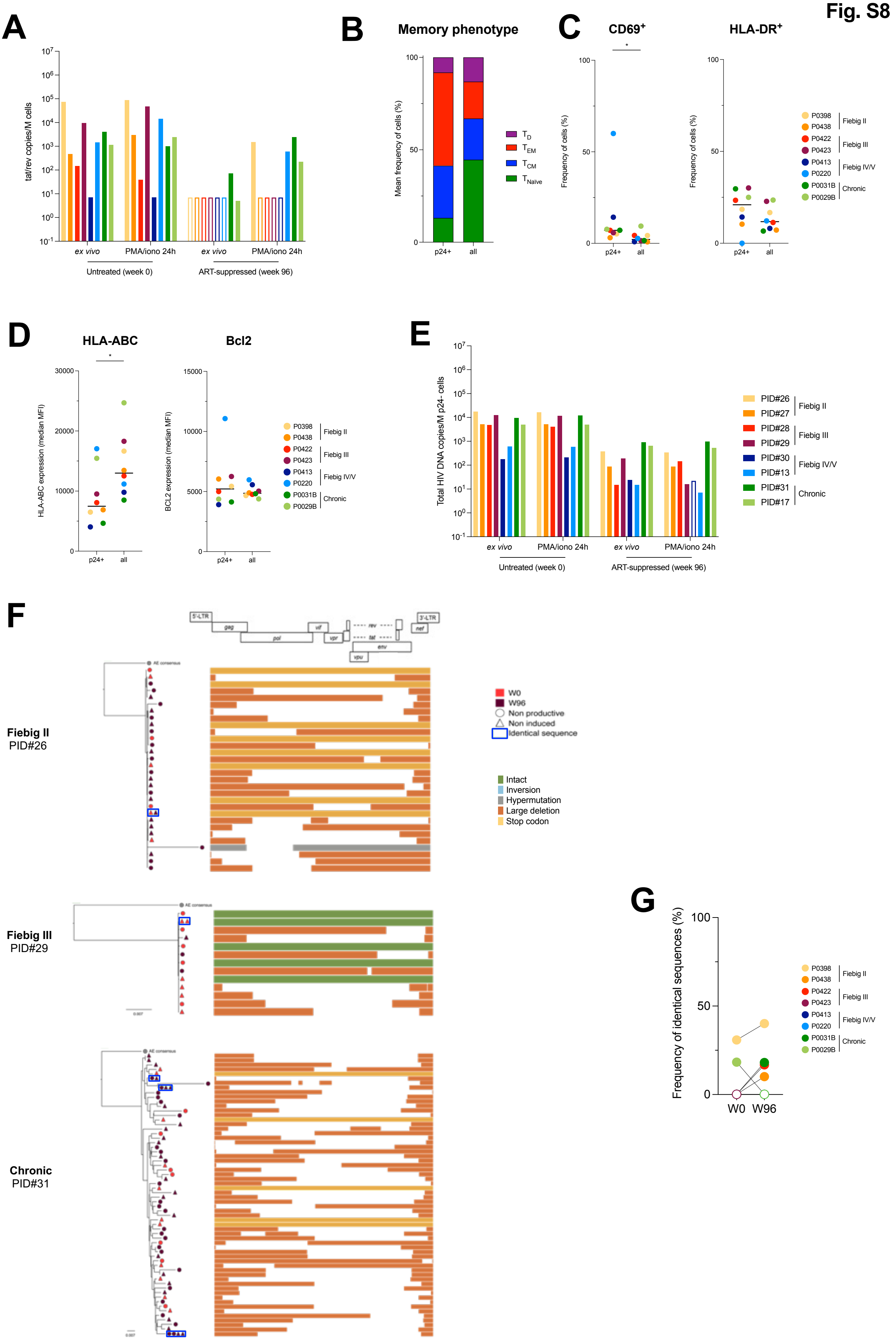
A
tat/rev copies/M cells
ex vivo
PMA/iono 24h
Untreated (week 0)
ART-suppressed (week 96)
B
Memory phenotype
Mean frequency of cells (%)
p24+
all
T D
T EM
T CM
T Naïve
C
CD69+
HLA-DR+
Frequency of cells (%)
P0398
P0438
P0422
P0423
P0413
P0220
P0031B
P0029B
Fiebig II
Fiebig III
Fiebig IV/V
Chronic
D
HLA-ABC
Bcl2
HLA-ABC expression (median MFI)
BCL2 expression (median MFI)
E
Total HIV DNA copies/M p24- cells
PID#26
PID#27
PID#28
PID#29
PID#30
PID#13
PID#31
PID#17
F
Fiebig II
PID#26
Fiebig III
PID#29
Chronic
PID#31
W0
W96
Non productive
Non induced
Identical sequence
Intact
Inversion
Hypermutation
Large deletion
Stop codon
G
Frequency of identical sequences (%)
W0
W96

**Fig. S8. Inducibility, intactness and persistence of latent proviruses in acute infection and on ART. A.** Total RNA from unstimulated or stimulated CD4+ T cells was used to quantify multiply spliced RNA (tat/rev). HIV transcripts were normalized to the number of input cells. **B.** Proportion of each memory subset in p24+ cells and total CD4+ T cells at the viremic time point *ex vivo*. Infected cells were overrepresented in the $T_{EM}$ subset. **C.** Activation status of infected cells. The frequency of each subset (CD69+, HLA-DR+) is depicted *ex vivo* for each participant according to the cell population (p24+ and total CD4+ T cells). **D.** Phenotype of infected cells. The median fluorescence intensity (MFI) for each marker (HLA-ABC, Bcl2) is depicted *ex vivo* for each participant according to the cell population (p24+ and total CD4+ T cells). **E.** p24- cells were bulk sorted for total HIV DNA quantification. Frequency of HIV DNA+ cells are presented in bar charts. Empty bars represent undetectable measures, and the limit of detection is plotted. **F.** Near-full length genome amplification was also performed on p24- cells to assess the genetic intactness of proviral genomes. To also test for inducibility, p24- cells from both ex vivo and stimulation conditions were included. Phylogenetic trees representing the proviral landscape of 3 participants are depicted. On the left of the graph, sequences are sorted according to their color for before (red) and during suppressive ART (burgundy); and symbol for both ex vivo (circle) and after stimulation (triangle), representing the non-productive and non-induced latent HIV reservoirs, respectively. Identical sequences are framed and represented on the same tree branch. On the right of the graph, proviral sequences are mapped on the HIV genome, and color-coded as follows: intact sequences in green, inversions in blue, hypermutations in grey, large deletion in orange and stop codons in yellow. **G.** The frequency of identical proviral sequences before and after ART in p24- cells is represented for each participant. (Wilcoxon; $p<0.05$, *; $p<0.01$**; $p<0.001$, ***; $p<0.0001$ ****).

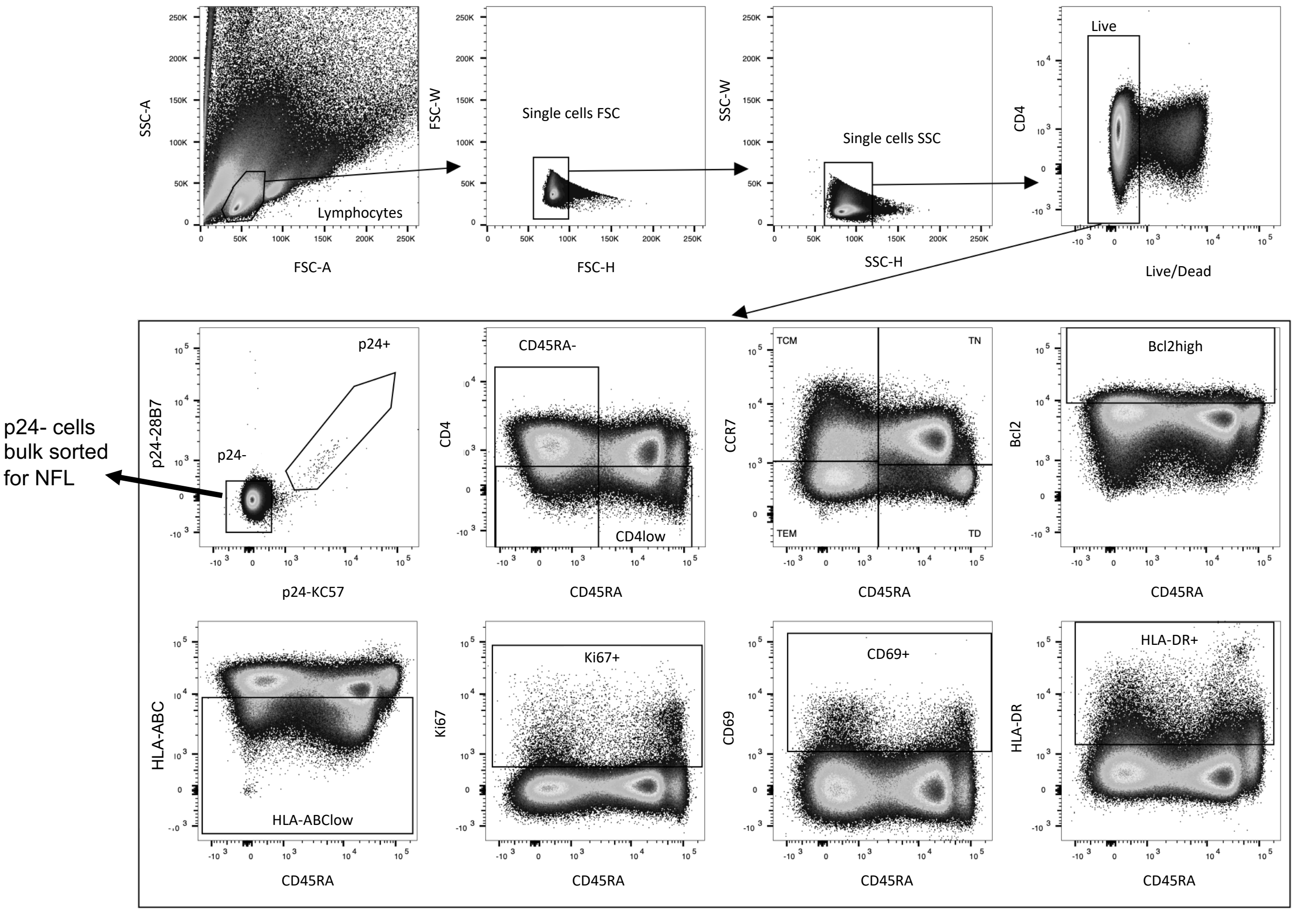


**Fig. S9. Gating strategy for near-full length HIV genome sequencing.** Gating strategy used to quantify *ex vivo* and induced p24+ cells before and after 96 weeks of ART. Bulk p24- cells were sorted for proviral HIV DNA quantification and near-full length sequencing (NFL).

### Table S1. Participants' characteristics

| PID# | Analysis | | | | Fiebig stage | Age at Week 0 (years) | Gender | HIV subtype | Week 0 (Viremic) | | | Week 96 (ART) | | |
|---|---|---|---|---|---|---|---|---|---|---|---|---|---|---|
| | **Pheno** | **TCR** | **Env** | **NFL** | | | | | CD4 (/mm3) | CD8 (/mm3) | HIV-RNA (copies/mL) | CD4 (/mm3) | CD8 (/mm3) | HIV-RNA (copies/mL) |
| PID01 | X | X | | | 1 | 28 | Male | CRF01_AE | 572 | 334 | 37142 | NA | NA | NA |
| PID02 | X | X | | | 1 | 25 | Male | CRF01_AE | 577 | 346 | 3385 | NA | NA | NA |
| PID03 | X | X | | | 1 | 26 | Male | CRF01_AE | 575 | 513 | 11969 | NA | NA | NA |
| PID04 | X | X | | | 2 | 27 | Male | CRF01_AE | 158 | 202 | 2024617 | NA | NA | NA |
| PID05 | X | X | | | 2 | 23 | Male | CRF01_AE | 283 | 226 | 6029465 | NA | NA | NA |
| PID06 | X | X | | | 2 | 26 | Male | CRF01_AE/B Recombinant | 278 | 319 | 6046140 | NA | NA | NA |
| PID07 | X | X | | | 3 | 22 | Male | CRF01_AE/B Recombinant | 256 | 352 | 6880660 | NA | NA | NA |
| PID08 | X | X | | | 3 | 25 | Male | CRF01_AE/B Recombinant | 509 | 1045 | 4631002 | NA | NA | NA |
| PID09 | X | X | | | 3 | 28 | Male | CRF01_AE | 656 | 820 | 4436997 | NA | NA | NA |
| PID10 | X | X | | | 4 | 25 | Male | CRF01_AE | 346 | 677 | 366196 | NA | NA | NA |
| PID11 | X | X | | | 4 | 34 | Male | CRF01_AE | 456 | 4556 | 6596826 | NA | NA | NA |
| PID12 | X | X | | | 4 | 21 | Male | B | 236 | 370 | 110572 | NA | NA | NA |
| PID13 | X | X | | | 5 | 40 | Male | CRF01_AE | 374 | 1869 | 686329 | NA | NA | NA |
| PID14 | X | X | | X | 5 | 39 | Male | CRF01_AE | 625 | 3936 | 277506 | 632 | 610 | <20 |
| PID15 | X | X | | | Chronic | 50 | Male | Not Done | 333 | 1941 | 21377 | NA | NA | NA |
| PID16 | X | X | | X | Chronic | 21 | Male | Not Done | 227 | 688 | 63588 | Not Done | Not Done | <34 |
| PID17 | X | X | | | Chronic | 26 | Male | Not Done | 616 | 1170 | 55844 | NA | NA | NA |
| PID18 | X | | X | | 1 | 24 | Male | CRF01_AE | 457 | 486 | 146194 | NA | NA | NA |
| PID19 | X | | X | | 1 | 25 | Male | CRF01_AE | 537 | 242 | 11674 | NA | NA | NA |
| PID20 | X | | X | | 2 | 23 | Male | CRF01_AE | 308 | 941 | 6010006 | NA | NA | NA |
| PID21 | X | | X | | 2 | 20 | Male | CRF01_AE | 307 | 297 | 68393 | NA | NA | NA |
| PID22 | X | | X | | 3 | 36 | Male | CRF01_AE | 488 | 3269 | 1107430 | NA | NA | NA |
| PID23 | X | | X | | 3 | 41 | Male | CRF01_AE/B Recombinant | 552 | 2036 | 7895089 | NA | NA | NA |
| PID24 | X | | X | | 5 | 20 | Male | CRF01_AE | 233 | 2531 | 5541200 | NA | NA | NA |
| PID25 | X | | X | | Chronic | 20 | Male | CRF01_AE | 345 | 865 | 19986 | NA | NA | NA |
| PID26 | | | | X | 2 | 26 | Male | CRF01_AE/B Recombinant | 249 | 430 | 736615 | 397 | 492 | <20 |
| PID27 | | | | X | 2 | 22 | Male | CRF01_AE | 641 | 481 | 11350800 | 1204 | 1058 | <20 |
| PID28 | | | | X | 3 | 22 | Male | CRF01_AE | 233 | 172 | 4416897 | 534 | 367 | <20 |
| PID29 | | | | X | 3 | 27 | Male | CRF01_AE/B Recombinant | 457 | 640 | 5286145 | 598 | 781 | <20 |
| PID30 | | | | X | 4 | 22 | Male | CRF01_AE | 571 | 552 | 14673 | 960 | 757 | <20 |
| PID31 | | | | X | Chronic | 45 | Male | CRF01_AE | 30 | 708 | 723346 | 267 | 1244 | <34 |

M, male; ART, antiretroviral therapy; TCR; T-cell receptor sequencing substudy; Env, HIV *Env* C2-V5 sequencing susbtudy; NFL, near-full length HIV genome sequencing susbtudy; NA, not applicable.

**Table S2. Primers used for amplification and sequencing of TCRβ, HIV *env* and near-full length HIV genomes**

| Name | Sequence 5'-3' |
|---|---|
| **PCR1 : Forward primers (tagged with M13F)** | |
| **VB2** | GTAAAACGACGGCCAGTATACTTCTATTGGTACAGACAAATCTTGG |
| **VB3** | GTAAAACGACGGCCAGTCTATGTATTGGTATAAACAGGACTCTAAG |
| **VB4** | GTAAAACGACGGCCAGTCAYARSGCTATGTATTGGTACAAGC |
| **VB5/9** | GTAAAACGACGGCCAGTCACTGTGTCCTGGTACCAACAG |
| **VB6** | GTAAAACGACGGCCAGTTACATGTACTGGTATCGACAAGACC |
| **VB7** | GTAAAACGACGGCCAGTTACCCTTTATTGGTACCGACAGAGCCTGG |
| **VB11** | GTAAAACGACGGCCAGTCTTTACTGGTACCGGCAGAWCYTGG |
| **VB12** | GTAAAACGACGGCCAGTTTTTCTGGTACAGACAGACCATGATG |
| **VB13** | GTAAAACGACGGCCAGTCACTGTCTACTGGTACCAGCAGG |
| **VB14** | GTAAAACGACGGCCAGTTGGACATGATAATCTTTATTGGTATCGAC |
| **VB15** | GTAAAACGACGGCCAGTCATGTACTGGTACCAGCAGAAGTC |
| **VB16** | GTAAAACGACGGCCAGTGTTATGTTTTTTGGTACCAACAGGTCC |
| **VB17** | GTAAAACGACGGCCAGTCATGTTTGTTCACTGGTACCGACAGAATC |
| **VB18** | GTAAAACGACGGCCAGTAGTCATGTTTACTGGTATCGGCAGC |
| **VB19** | GTAAAACGACGGCCAGTTGCCATGTACTGGTACCGACAG |
| **VB20** | GTAAAACGACGGCCAGTCACAACTATGTTTTGGTATCGTCAG |
| **VB21** | GTAAAACGACGGCCAGTTAGTTATGTTTACTGGTATCATAAGACGC |
| **VB23** | GTAAAACGACGGCCAGTATACTTTTTGTTTATTGGTATCAACAGAATCAG |
| **VB24** | GTAAAACGACGGCCAGTATGTACTGGTATCGACAAGACCC |
| **VB25** | GTAAAACGACGGCCAGTTGACAAAATGTACTGGTATCAACAAGATC |
| **VB29** | GTAAAACGACGGCCAGTTGATGTTCTGGTACCGTCAGCAAC |
| **VB30** | GTAAAACGACGGCCAGTCAACCTATACTGGTACCGACAGG |
| **PCR1 : Reverse primers (tagged with M13R)** | |
| **JB1-1** | CAGGAAACAGCTATGACCAACTGTGAGTCTGGTGCCTTGTCCAAAG |
| **JB1-2** | CAGGAAACAGCTATGACAACCTGGTCCCCGAACCGAAGG |
| **JB1-3** | CAGGAAACAGCTATGACAACAGTGAGCCAACTTCCCTCTCCAAAATA |
| **JB1-4** | CAGGAAACAGCTATGACCAGAGAGCTGGGTTCCACTGCCAAAAAACA |
| **JB1-5** | CAGGAAACAGCTATGACAGAGTCGAGTCCCATCACCAAAATGC |
| **JB1-6** | CAGGAAACAGCTATGACCTGGTCCCATTCCCAAAGTGGAGG |
| **JB2-1** | CAGGAAACAGCTATGACAGCCGTGTCCCTGGCCCGAAGAAC |
| **JB2-2** | CAGGAAACAGCTATGACCGTTTTTTGGAGAAGGCTCTAGGCTGACC |
| **JB2-3** | CAGGAAACAGCTATGACCAGCCGGGTGCCTGAGCCAAAATAC |
| **JB2-4** | CAGGAAACAGCTATGACCGGGTCACGGCGCCGAAGTAC |
| **JB2-5** | CAGGAAACAGCTATGACAGCCGCGTCCTGGCCCGAAG |
| **JB2-6** | CAGGAAACAGCTATGACCTGCCGGCCCCGAAAGTCAGG |
| **JB2-7** | CAGGAAACAGCTATGACCCTGGTGCCCGACCCGAAG |
| **PCR2 and Sequencing: Forward and Reverse primers** | |
| **M13F** | GTAAAACGACGGCCAGT |
| **M13R** | CAGGAAACAGCTATGAC |
| **PCR3 (MiSeq adaptors) : Forward and Reverse primers** | |
| **CS1-M13F** | ACACTGACGACATGGTTCTACAGTAAAACGACGGCCAGT |
| **CS2-M13R** | TACGGTAGCAGAGACTTGGTCTCAGGAAACAGCTATGAC |
| **PCR1 : HIV Env AE C2-V5** | |
| **Env7** | AATGGCAGTCTAGCAGAAG |
| **OutV5R_AE** | TCCTAGTGGTTCAATTTGTACTAC |
| **PCR2 and Sequencing : HIV Env AE C2-V5** | |
| **Env7** | AATGGCAGTCTAGCAGAAG |
| **InV5R_AE** | ACTTCTCCAATTGTCCTTTA |
| **PCR1: near full-length HIV AE** | |
| **263-AE-F** | AGGGACTCGAAAGCGRAAGT |
| **BlouterR** | TGAGGGATCTCTAGTTACCAGAGTC |
| **PCR2: near full-length HIV AE** | |
| **652-AE-F** | ACTCGAAAGCGRAAGTTCCAGAG |
| **280-AE-R** | CTAGTTACCAGAGTCCTAACACAGAYG |
